# CellTune: An integrative software for accurate cell classification in spatial proteomics

**DOI:** 10.1101/2025.05.05.652215

**Authors:** Yuval Bussi, Dana Shainshein, Eli Ovits, Sarah Posner, Nofar Azulay, Noa Maimon, Tal Keidar Haran, Raz Ben-Uri, Caitlin Brown, Noam Schuldiner, Eylon Yaniv, David Van Valen, Idan Milo, Ofer Elhanani, Robert Schiemann, Leeat Keren

## Abstract

Spatial proteomics measures multiple proteins *in situ*, capturing tissue complexity. However, cell classification in densely packed tissues remains challenging due to the lack of efficient classification algorithms, annotation tools, and high-quality labeled datasets to benchmark computational methods. We introduce *CellTune*, an integrated software for analysis of large spatial proteomics datasets, which streamlines precise cell classification through an optimized human-in-the-loop active learning workflow. It advances core capabilities across within a unified, intuitive, and code-free interface. To evaluate CellTune, we created *CellTuneDepot*, a resource of 40k manually-annotated cells and 3.5 million high-quality labeled cells across 60 cell types. CellTune outperforms alternative methods, achieving accuracy comparable to human performance while enabling increased classification resolution and discovery of novel cell types. Together, CellTune and CellTuneDepot provide researchers with a tool for state-of-the-art classification accuracy and resolution at scale to drive biological insights.

## Introduction

The spatial organization of tissues facilitates healthy function, and its disruption contributes to disease^1^. Spatial proteomics enables the measurement of multiple proteins *in situ*, offering unprecedented insights into the molecular and spatial complexity of tissues. While rapid technological advancements in spatial proteomics have sparked considerable interest in the scientific community^2–14^, data analysis remains a major obstacle to broader adoption. Specifically, transforming protein expression images into structured maps of segmented and classified cells is an inaccurate and laborious process, requiring researchers to navigate fragmented pipelines and visualization software.

The analysis of multiplexed imaging data typically follows a structured yet disjoint pipeline. First, cells are segmented into tabular data structures, with protein expression quantified for each cell. This table serves as input for classification, where cell types and phenotypes are inferred based on co-expressed proteins and prior knowledge. Early methods for cell classification relied on manual gating or clustering of the expression table using algorithms that were developed for measurements of cells in suspension, such as cytometry or single cell RNA sequencing (scRNAseq) ^5,10,15–22^.

However, deriving cell classifications from multiplexed images has unique challenges over classifying cells in suspension. First, the densely packed nature of tissues, coupled with irregular cell shapes, results in spillover of protein signals onto neighboring cells and complicates accurate segmentation and classification. This is exacerbated by the two-dimensional imaging of inherently three-dimensional tissue structures, which can obscure important spatial information^23^. Second, datasets are highly imbalanced, with rare cell populations often defined by subtle expression patterns of just one or two markers, making them difficult to identify using conventional methods. Third, technical variability, including differences in tissue quality, imaging artifacts, and antibody performance, adds another layer of complexity, necessitating tools that are both precise and adaptable.

Approaches to deal with the challenges of cell classification in multiplexed images have focused on incorporating spatial information in a myriad of ways, including compensation^24^, pixel clustering^25^, graph-based modelling^26,27^ and image-based deep learning ^28,29^. While advantageous, spatial information is not sufficient, and to discern real and spurious coexpression patterns it is necessary to infuse prior knowledge. Since most approaches ultimately perform unsupervised clustering, the integration of prior knowledge is usually performed post-hoc by visual inspection and manual manipulation and annotation of the clustering results. Although beneficial in allowing the researcher to identify novel cell types, this process is labor-intensive and time-consuming and does not enhance the algorithm for subsequent analyses. In contrast, alternative approaches infuse prior knowledge before classification using predefined cell types ^26,30^, or labeled examples for machine learning ^31,32^ or deep learning ^27–29^. Importantly, to realize their promise and avoid being confined to a limited number of predefined cell types, these methods require accurate training data, drawn from the same distribution as the cells for classification. Crucially, such ground-truth labels are lacking.

The lack of ground truth labels for cell classification in spatial proteomics presents a major bottleneck in the field. Accurate labels are needed for both training and evaluating new classification algorithms. However, generating high-quality training data suffers from the same issues described above, and therefore necessitates extensive manual curation. This stands in contrast to segmentation, where the availability of large-scale labeled datasets has already enabled the development of accurate and robust segmentation models ^33–36^. Moreover, while software for visualization and analysis of multiplexed images is being developed ^37–39^, the tools available for manual labeling are not well-adapted for multiplexed images. Performing tasks such as moving between cells, visualizing the appropriate proteins, and assigning cell types is cumbersome and slow. As a result, existing manually-labeled resources are limited to easily classifiable cells, small image subsets, or few cell types ^29,40,41^, and a comprehensive dataset of ground truth manually-curated cell classifications to serve as a benchmark for method development is lacking.

To address these challenges, we developed CellTune, an integrated software platform for visualizing, annotating, and classifying multiplexed tissue images. CellTune combines spatial feature computation with a human-in-the-loop active learning framework, enabling iterative refinement of classification models guided by expert input. This approach strikes a balance between leveraging prior biological knowledge and maintaining flexibility for exploratory discovery, significantly enhancing the accuracy and efficiency of cell classification. Uniting analysis and visualization into a single accessible platform, CellTune is specifically designed to handle large spatial proteomics datasets, encompassing hundreds of multiplexed images. The platform simplifies tasks such as cell classification by automatically displaying relevant combinations of protein channels, facilitating rapid navigation across images, and allowing simultaneous interaction with protein images and cell populations across an entire cohort.

To support the development and benchmarking of CellTune, we created CellTuneDepot, the largest manually annotated dataset for spatial proteomics to date, featuring over 40,000 manually labeled cells from multiple tissue types, alongside millions of high quality CellTune-generated labels. Benchmarking CellTune against 10 state-of-the-art methods demonstrated its superior performance, with a 10–15% improvement in accuracy, achieving human-level performance. CellTune doubled the number of distinguishable cell types, enabling finer-grained classification and facilitating the discovery of new cell populations by interactively engaging users with ambiguous classifications. Together, CellTune and CellTuneDepot address critical limitations in current analysis workflows, offering a robust, scalable, and biologically informed framework for cell classification.

## Results

### CellTune: a human-in-the-loop approach for cell classification

We designed CellTune as an active learning pipeline for cell classification in multiplexed images, guided by the following principles: (1) combining analysis and visualization into a single software, (2) learning from human knowledge, (3) utilizing spatial features in classification, and (4) minimizing human curation efforts without compromising data quality. The classification pipeline is designed as a series of iterative cycles, each involving training a classification model, prioritizing cells for manual annotation and efficient labeling of a subset of cells. This human-in-the-loop approach gradually improves the model’s performance in cell classification (**Fig. 1A-D, S1A**).

**Figure 1.**
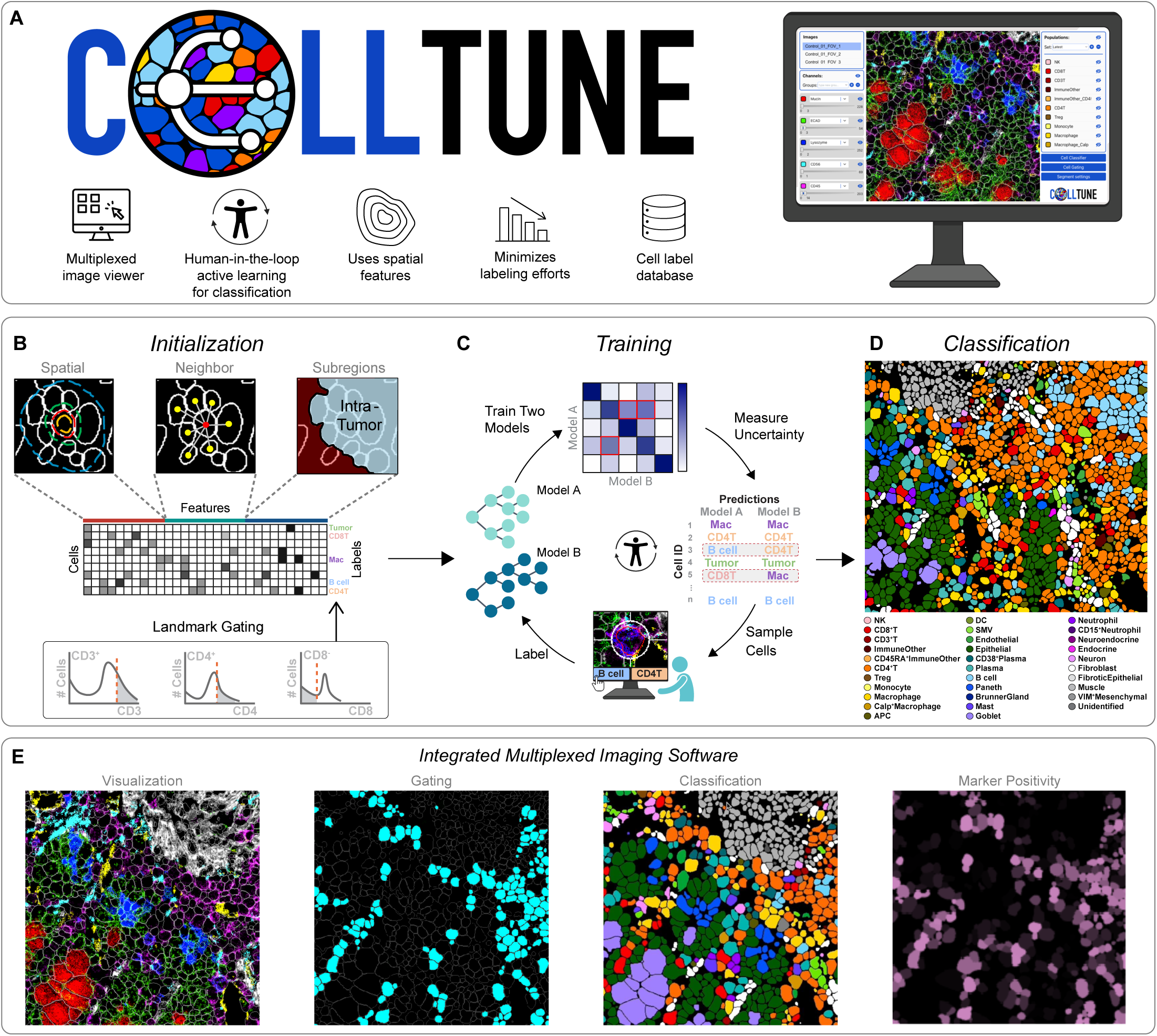
Overview of CellTune. **(A)** Schematic overview of CellTune. The graphical user interface is shown on the right, illustrating the image viewer with segmentation overlays and interactive panels for images, channels, and populations. **(B)** Initialization of the cell classification pipeline begins with feature extraction, including protein expression per cell, spatial and neighborhood features, and very stringent gating of a small set of easy-to-classify landmark cells. **(C)** Active learning-based training and classification pipeline: two gradient-boosted tree models are trained in parallel. Their results are compared to identify areas of uncertainty, and cells from these regions are prioritized for human review. The updated labels are iteratively used to improve the model. **(D)** Final cell type classification. **(E)** CellTune’s key features, including multi-channel image visualization, cell gating, cell classification, and marker positivity prediction.

To initialize CellTune, we input multiplexed images and their corresponding segmentations, which can be generated with various segmentation algorithms ^33–35^. We then compute a variety of features on each cell (**Fig. 1B, S1B**). Importantly, these features incorporate spatial information, including the expression of each protein in varying erosions and expansions of the cell’s segmentation mask, in neighboring cells, and other spatial features, such as whether the cell resides within a specific anatomical structure (e.g., tumor, vessel, or muscle). Next, we perform very stringent gating of *landmark* cells, which can be rapidly classified with high confidence according to prior knowledge (**Fig. 1B**) ^42^. For example, a cell that expresses strong CD8 and CD3 and no other lineage protein is a landmark CD8^+^ T cell. Exploratory clustering and visualization can supplement prior knowledge and further assist with identifying any prevalent dataset-specific cell types that should be landmarked (**Fig. S1C-D**). In our experience, this landmarking process enables to quickly and easily classify a minority of the data (between 10-20% of the cells). This data serves as input to train an initial classifier (**Fig. 1C, S1E**). Since this classifier was only trained on cells that are easy to classify, it did not encounter cells spanning the real-world distribution of protein expression in the images and therefore has low accuracy (40-70% depending on the dataset). It is also limited to cell types anticipated by the researcher.

Following this initialization, we continue to cycles of model refinement using a query-by-committee (QBC) framework^43^ (**Fig. 1C, S1E**). In each cycle, two models – e.g. XGBoost^44^ and CatBoost^45^– are trained on the labeled data that has accumulated so far, and their agreement is evaluated on new, unlabeled cells. The premise is that easy cells and cells which are similar to those already labeled, are very likely to be classified the same, and correctly, by both models. In contrast, cells classified differently by the models are enriched for more challenging cells or novel cell types. These cells present good candidates for manual labeling, since they encompass novel information and adding them to the training data holds high potential for model-improvement. Both XGBoost and CatBoost are gradient-boosted decision tree (GBDT) models, which are particularly well-suited for tabular data and effective under limited label availability, as is typical in spatial proteomics. Their inherent feature selection helps mitigate overfitting, while their distinct tree-building strategies ensure that the models learn differently – an essential property for the QBC framework which relies on model disagreement to identify informative cells for labeling.

Next, CellTune automatically samples differentially classified cells for labeling, with the number of sampled cells adjustable by the user (**Fig. 1C, S1E**). This automated selection process strives to minimize manual labeling and focus effort on cells that are most likely to improve model performance by balancing model agreement across cell types and images, while also enriching for rare cell types. Users can optionally customize the sampling to emphasize specific cell types or confusions of interest. The selected cells are then presented to the researcher in an efficient manner that expedites labeling. Following labeling, manually labeled cells are added to the training data, for another round of model training, prediction, and evaluation. These cycles are repeated until the desired accuracy level is reached and then classifications are finalized (**Fig. 1D**).

Altogether, CellTune combines several machine learning strategies, including human-in-the-loop, query-by-committee, and prioritized sampling to direct human annotation efforts, with the goal of expediting the training of a data-relevant cell classification model. A complete description of each one of these steps is detailed in the methods section and supplementary Figure S1. Figure S2 provides snapshots of how these steps are implemented in the CellTune software.

### CellTune provides a multiplexed image viewer with cell labeling functionality

The CellTune workflow is implemented in an integrated desktop application that displays multiplexed images, together with various views including cell segmentation and classification (**Fig. 1E, 2A**). CellTune was specifically designed for viewing and analyzing large cohorts of multi-channel pathology imaging. Unlike many software, which are image-centric, CellTune works on a project, which can include hundreds of images, each with multiple channels. It rapidly transitions between images, while preserving the visualization settings, including which channels are on, their histogram scaling, the segmentation, and populations. Cell populations are shared across the entire cohort, and populations can be easily created, merged, or partitioned. To minimize the time spent on selecting and adjusting channels for visualization, it allows the user to easily define *channel groups*, pre-defined sets of channels that can be visualized together to explore specific groups of cells (**Fig. 2B**). Gating is performed using a dedicated widget in the software, in which the researchers can define complex logical gates and interactively visualize the gated cells together with the relevant channels across different images, allowing them to adjust thresholds in real time (**Fig. 2C**). This enables exploration and refinement of cell populations, as well as the evaluation of classification results.

**Figure 2.**
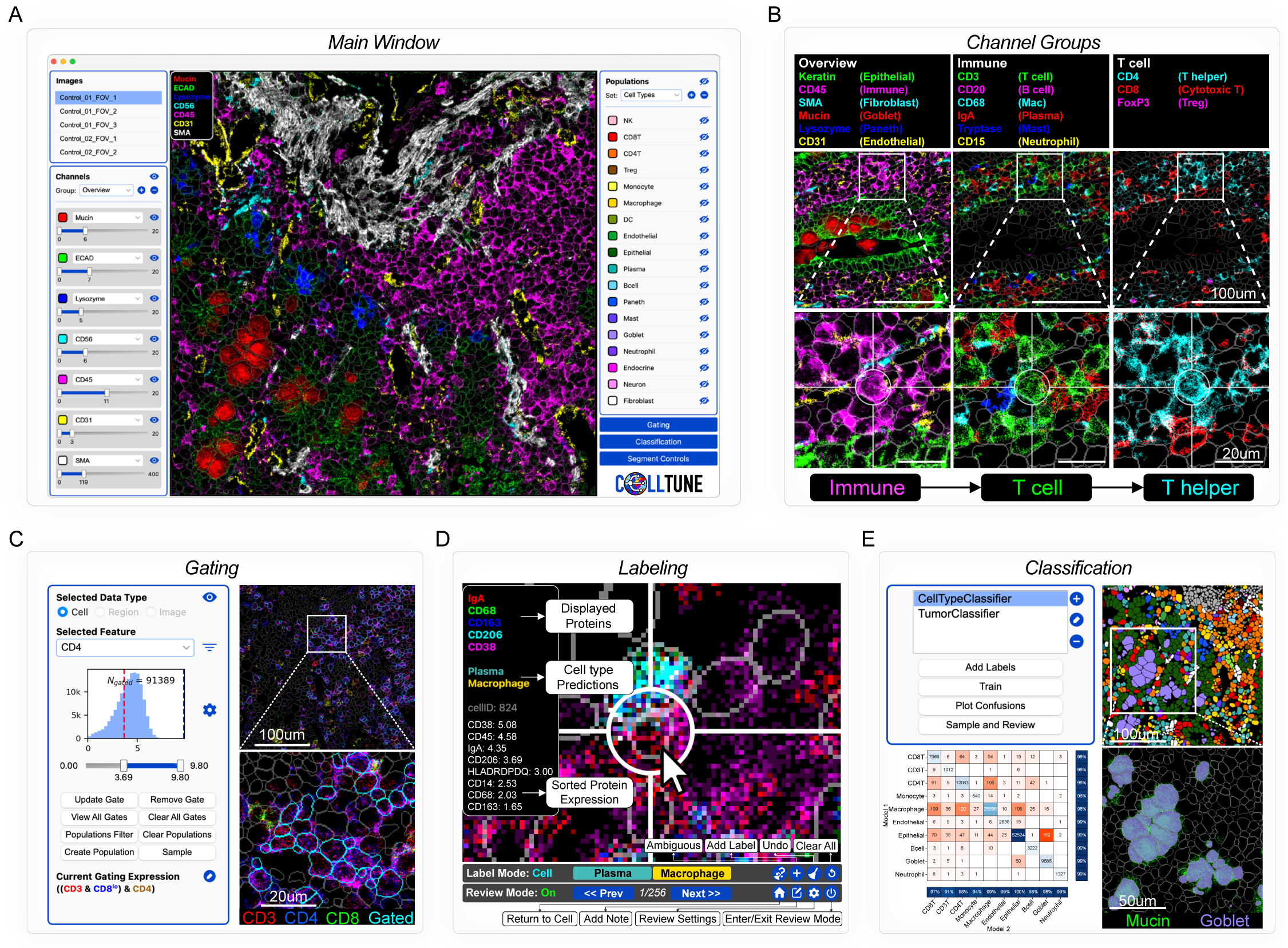
CellTune integrates visualization, gating, labeling, and classification in a software optimized for analyzing highly multiplexed images. **(A)** Main window of the CellTune software. The central image viewer is flanked by image and channel selection controls on the left and population, classification, gating, and segment controls on the right. (**B)** CellTune enables the user to define groups of channels that are often visualized together, along with their display settings, for rapid informative visualization. **(C)** Cell gating supports complex expression rules, such as identifying cells strictly positive for CD3 and CD4 while negative for CD8. **(D)** CellTune’s interactive labeling mode streamlines the labeling process by automatically adjusting the zoom, displayed proteins, providing cell information on hover, and presenting a labeling and review panel. **(E)** Top left: Classification control panel. Bottom left: The user can display a confusion matrix between the predictions of the two trained models. Top right: Cells are colored by classification. Bottom right: The user can evaluate the classification by overlaying protein expression (e.g., Mucin, shown in green) and classified cell types (e.g., Goblet, shown in red).

A key step in the CellTune workflow is manual curation to generate ground-truth labels for model training. During this process, cells are presented to the user alongside conflicting cell-type predictions from the models. The user must then visually inspect the cells and decide whether one of the labels is correct, both are equally plausible, or neither is accurate. Additionally, the user may define a novel class if needed (**Fig. 2D**). This correction process, while faster than labeling from scratch, remains challenging and time-consuming. To manually label cells in multiplexed images, researchers must examine the expression of several proteins within the target cell. However, simultaneously visualizing dozens of proteins through color overlays is impractical, as colors quickly overlap. Consequently, a significant portion of the time spent on manual labeling is devoted to adjusting the displayed proteins and their corresponding colors.

To streamline the labeling process, CellTune incorporates several innovations. Cells are presented sequentially, with the interface automatically zooming in on the cell of interest. This facilitates efficient navigation and ensures a focused view of each cell. In addition, the software automatically displays the channels most relevant for resolving a specific confusion, together with those most strongly expressed in the specific cell. For example, if a cell is ambiguously classified as either Plasma or Macrophage, the channels associated with these cell types (IgA, CD38, CD68, CD163, CD206) are automatically displayed when the cell is selected (**Fig. 2D**). Researchers can still customize the displayed channels as needed. Cells requiring labeling are also organized by the suggested cell-type pairs predicted by the models. Since each cell type is linked to a specific set of proteins, this organization reduces the time and cognitive effort required to adapt to new channels. Furthermore, when hovering over a cell, CellTune displays several key details: the currently visible proteins, the model predictions, and a ranked list of protein expression values for that cell, from strongest to weakest (**Fig. 2D**). This feature enables researchers to quickly determine whether additional proteins should be visualized.

Altogether, the process is designed to improve the accuracy and speed of cell classification. The software enables smooth transitions between running the classification models, labeling and various views to evaluate the results (**Fig. 2E**). The researcher can decide to stop training at any point according to the level of accuracy that is desired for the study. The manually labeled cells and models are saved, such that the same researcher or another can resume the process at any point in time if new images are acquired and require further training, mistakes are identified in downstream analyses or a higher level of accuracy is required.

### CellTune cell classification achieves accurate human level performance

We used CellTune to label over 3.5 M cells across 5 datasets spanning different imaging technologies, organs, and pathologies **(Fig. 3A)**, classifying 22-32 cell types for each dataset, and over 60 cell types in all datasets. The pipeline took ∼80 hours on average, with ∼5000 cells manually labeled per dataset. Visual inspection showed good qualitative agreement between protein images and classified cell types, even in cases of closely-related or closely-packed cells **(Fig. 3B)**.

**Figure 3.**
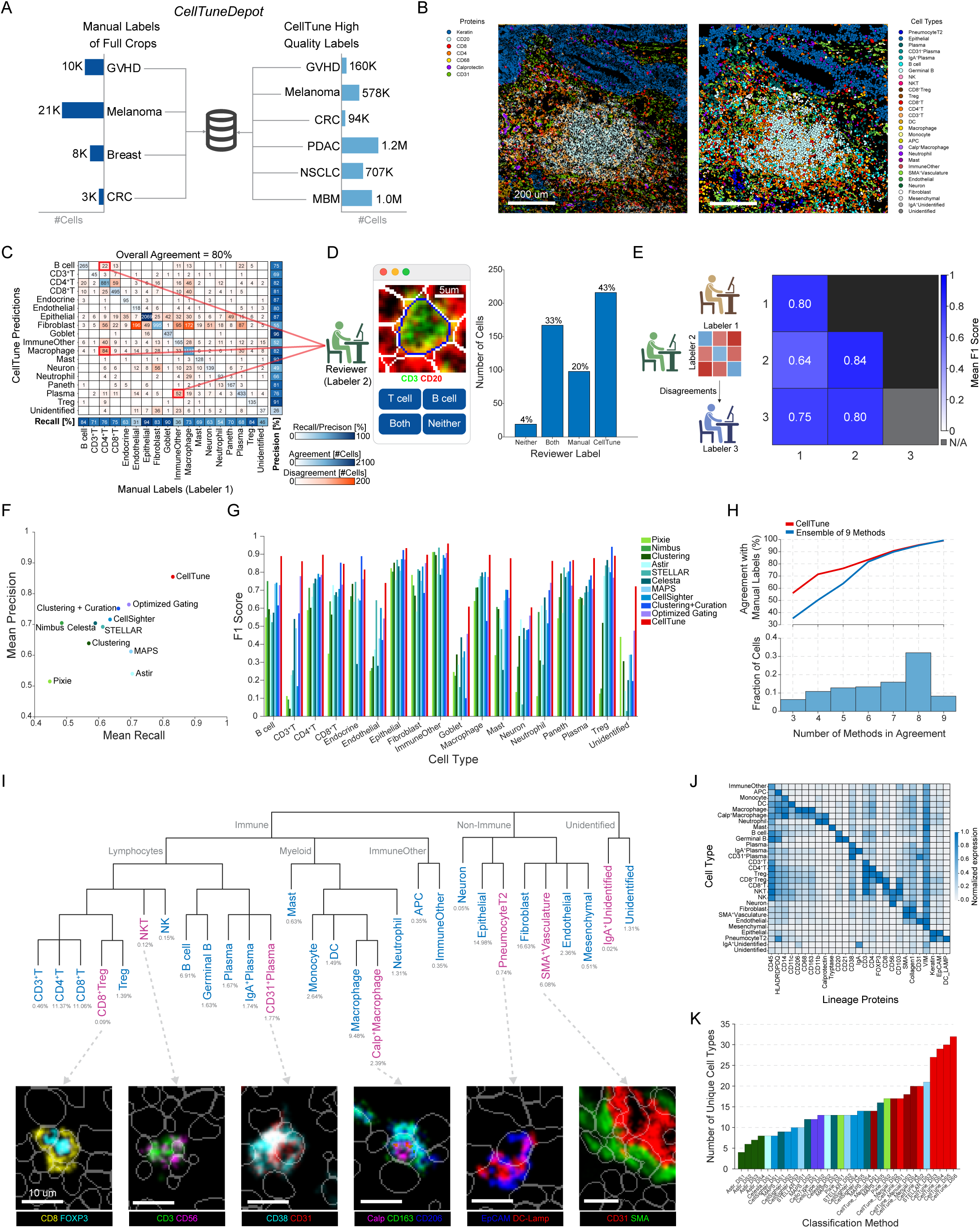
CellTune surpasses human accuracy, expands cell type classification, and enables large-scale high-quality labeling. **(A)** *CellTuneDepot*, a repository of manual labels of full crops and high-quality labels of entire images generated with CellTune. GVHD: graft-versus-host disease. CRC: colorectal cancer. PDAC: pancreatic ductal adenocarcinoma. NSCLC: non-small cell lung cancer. MBM: melanoma brain metastases. **(B)** An example field-of-view (FOV) from the NSCLC dataset displaying protein expression levels (left) alongside CellTune classifications (right). **(C)** The confusion matrix comparing between a manual labeler (x-axis) and CellTune (y-axis) on the GVHD dataset shows high agreement. **(D)** Manual labeling of 500 disagreeing labels by a different labeler (reviewer) shows higher agreement with CellTune. **(E)** Left: schematic for generating gold-standard consensus manual labels. Two manual labelers label the same cells. An expert third labeler, labels the cells where they disagreed, reviews their agreements, and curates the labels. (Right) Mean F1 agreement between different labelers (off-diagonal) and from each labeler to the consensus (diagonal). Comparing labeler #3 to the consensus was not applicable. **(F-G)** Comparison of CellTune to other cell classification methods by evaluating on the consensus labels generated in (E). Methods include: Pixie^25^, Astir^30^, Celesta^26^, MAPS^31^, STELLAR^27^, CellSighter^28^, and Nimbus^40^, clustering^22^, pixel clustering with manual corrections, and optimized gating. Where possible, methods were optimized using the consensus labels to give an upper bound of their best possible performance – for clustering, each cluster was given the optimal label and for optimized gating, thresholds were specifically adjusted to maximize performance (Methods). **(F)** Scatterplot of the mean recall (x-axis) vs mean precision (y-axis) for each method. **(G)** F1 scores for individual cell types. **(H)** Line plot comparing CellTune’s accuracy to an ensemble of nine methods. Agreement with manual labels (y-axis) is shown as a function of the number of methods agreeing on a classification in the ensemble (x-axis). The distribution of cells across agreement levels is shown as a histogram. For cells on which many methods agree, both methods converge to near-perfect agreement, while at lower consensus levels, CellTune maintains higher accuracy than the ensemble. **(I)** Top: Lineage tree of cell types classified in the NSCLC dataset using CellTune, with cell type frequencies displayed below each label. Magenta highlights cell types that were not initially gated by the researcher and were discovered following the CellTune pipeline. Bottom: Example images. **(J)** Mean expression of each lineage protein (x-axis) for each cell type (y-axis) in the NSCLC dataset. **(K)** Number of cell types classified in different datasets (**Supplementary Table 1)**. For each method, shown are the number of cell types reported on different datasets in the respective publication of the method. Using CellTune (red), users classified more cell types than other methods, and more than were manually labeled on the same datasets (maroon).

To quantitatively test the approach, we generated a comprehensive dataset of manual labels. Previous attempts to generate such data have been constrained in various ways: some focused only on ‘easy cells’ that are straightforward to gate, failing to capture real-world data distribution, while others were limited in the number of images, channels, or cell types classified^29,40^. Many publications have evaluated the performance of their cell classification algorithms on labels that were generated by applying a backbone algorithm + curation^27,28,31^. However, curating algorithm-generated labels strongly biases the results of the new model towards the original algorithm. To generate true gold-standard labels, we took an approach of randomly selecting crops from different images, each encompassing ∼200 cells, and manually labeling all the cells in the crop (Methods). Overall, we labeled >40K cells from four different datasets, which to our knowledge constitutes the largest fully manually labeled dataset of highly multiplexed images to date **(Fig. 3A)**.

We compared the performance of CellTune to the manual labels and found 80% agreement (**Fig. 3C**), which is on par with the concordance between two different human labelers^28,32^. The disagreement between CellTune and the manual labeler could reflect biases in the model, user-specific biases, or ambiguous cells, which can have two plausible labels. To further evaluate these results, we had a second manual labeler (Labeler 2) review and evaluate 500 cells whose classification differed between CellTune and Labeler 1. Labeler 2 was presented with two optional labels and had to decide whether they agreed with either one, none or both. The order of the labels was randomized, and Labeler 2 was blinded to which label was suggested by the model or by Labeler 1. We found that, according to Labeler 2, CellTune gave the correct classification in 43% of the cases, Labeler 1 in 20% of the cases, both were equally plausible in 33% of the cases and none were correct in 4% of the cases **(Fig. 3D)**.

This analysis, clearly favoring CellTune (Fisher’s exact test OR=2.9, p-value<10^14^), demonstrates two important points that should be considered when evaluating classification algorithms for spatial proteomics. First, the higher error rate of Labeler 1 over CellTune suggests that even gold-standard manual labels have a high error rate, since manually labeling comprehensive fields of view (FOVs) is challenging. A labeler may get tired or may not be sufficiently diligent to consider all relevant channels and labels. In contrast, CellTune’s human-in-the-loop approach points the researcher to plausible cell types to consider, which may reduce “labeling fatigue” and ultimately improve accuracy and speed. As such, CellTune classifications may be considered a better gold standard than manual labels. Second, this analysis suggests that a non-negligeable fraction of cells are ambiguous and could potentially have several labels. Dealing with such cases is outside the current scope of CellTune but presents interesting avenues for follow-up work.

To further evaluate CellTune’s performance, we compared it to other published methods. To improve the manual labels for this task, we had 10,000 cells from the graft-versus-host disease (GVHD) dataset labeled by two additional labelers. Cells which differed in their classification between labelers 1 and 2 were labeled by a third labeler, to generate a consensus label (**Fig. 3E**). Notably, these three labelers were independent from the labeler who generated the training data for this dataset during the human-in-the-loop annotation process. We then compared the agreement of these consensus labels to various methods including Pixie^25^, Astir^30^, Celesta^26^, MAPS^31^, STELLAR^27^, CellSighter^28^, and Nimbus^40^, as well as pixel clustering with manual corrections. For each method, we established an upper bound on performance by optimizing its running regime, parameter selection and training data where applicable to obtain the best results possible (Methods). Importantly, we provided 5,000 high-quality consensus labels for training supervised methods or parameter tuning of unsupervised methods – a resource that users typically wouldn’t have access to for their specific datasets without significant manual annotation effort. The remaining 5,000 consensus labels were reserved for evaluation. We also included clustering^22^, where each cluster was given the label that maximized the agreement with the ground truth, and optimized gating, where thresholds were specifically adjusted to maximize performance, in order to establish an upper bound for these two methods, which are among the most commonly used for cell classification. We found that CellTune achieved the highest accuracy for the entire dataset, with an F1 of 0.84, compared to F1 scores of 0.45-0.71 for the other methods (**Fig. 3F,G**). All methods showed variability between the accuracy of classification of different cell types, with some cell types being consistently easier to classify across methods, such as epithelial cells or Tregs, and others being consistently more difficult, such as endothelial cells or neurons, which are more difficult to segment in 2D and are often part of tissue structures. Unidentified cells, defined as negative for all lineage markers, were also challenging to classify for all methods. For all cell types, CellTune outperformed the other methods **(Fig. 3G)**, except for Tregs and endocrine cells, where only manual curation achieved slightly higher accuracy (Tregs 0.89 vs. 0.94, endocrine (0.81 vs 0.88). These exceptions likely reflect the fact that these two rare cell types are defined by very specific, easily curated markers (FOXP3 and Chromogranin-A, Methods). For easier cells, on which more than six methods agreed on their classification, CellTune performed as well as the majority vote ensemble of all other methods (Methods). For more difficult cells, on which 3-5 methods agreed, CellTune demonstrated an improvement of 15-20% in classification accuracy relative to the majority vote of all other methods (**Fig. 3H**). Overall, CellTune demonstrates increased accuracy relative to existing approaches, as evaluated using a gold-standard manually-labeled dataset.

### CellTune naturally incorporates supervised and unsupervised learning for increased resolution and identification of new cell types

One prominent advantage of the human-in-the-loop approach is that often the data contains unexpected cell types, defined by novel combinations of protein expression that were not necessarily anticipated in the study design. For example, in a panel that we designed to study lung cancer, we included pan-Keratin as a lineage protein of epithelial cells and LAMP-3 as a protein expressed in a subset of dendritic cells (DCs) ^46^. Running CellTune on this dataset brought up many confusions between epithelial cells and DCs. Manual inspection of these cells indicated that the cells morphologically appeared as epithelial cells and they were indeed expressing both pan-Keratin and LAMP-3 **(Fig. 3I)**. Inspection of the literature indicated that these are type-2 pneumocytes ^47^, known to express LAMP-3. The CellTune pipeline easily allowed us to gate a landmark population of Type-2 Pneumocytes, add it to the training data and retrain CellTune with this new label. Similarly, CellTune presented confusions between ‘cytotoxic T cells’ and ‘T regulatory cells’, pointing to the existence of FOXP3^+^ CD8^+^ T cells ^48^ in this dataset, as well as confusions between ‘cytotoxic T cells’ and ‘NK cells’, pointing to the existence of CD56^+^ CD8^+^ NKT cells^49^ **(Fig. 3I).** Notably, CellTune easily surfaced these cells, even though their abundance in the dataset vas very low (0.09% and 0.12% respectively). These examples showcase CellTune’s flexibility and ability to consolidate prior knowledge with novel findings.

We hypothesized that CellTune’s ability to flag unexpected cell types will increase the overall number of cell types in a dataset and allow the detection of rare cell types. Indeed, when comparing CellTune labels to the initial manual gates for the lung cancer dataset, we found that using CellTune the researcher was able to detect 7 new cell types, even cells with low frequencies of <0.1% **(Fig. 3I,J)**. To further explore this property, we compared the number of cell types detected by CellTune to the number of cell types detected by other methods. Many of the methods ultimately require clustering, and the number of clusters can be subjectively altered by the researcher. Therefore, to avoid biasing the results to our selection of parameters, for each method we relay the number of cell types reported by the authors in their original publication. The number of potentially detectable cell types depends on the tissue and marker panels, so we restricted this analysis to studies that imaged a few dozen proteins, and should theoretically yield similar numbers of cell types (Methods). We found that CellTune consistently produced more and described finer-grain cell types, even where the same dataset was used, with a range of 17-32 (mean = 25.8) compared to 4-21 (mean = 11.4) for all others (**Fig. 3K, Supplementary Table 1**). This highlights CellTune’s ability to resolve complex and rare cell types at a resolution that other methods may theoretically achieve but rarely reach in practice due to limitations in their workflows and annotation tools. CellTune also produced more cell types than the number of cell types generated by manual labeling, and detected more cell types with low frequency than other methods (**Fig. 3C,G**). Altogether, CellTune combines supervised and unsupervised learning for increased resolution and identification of new and infrequent cell types.

### CellTuneDepot expedites and improves analysis of novel datasets

Applying CellTune to analyze six different datasets resulted in CellTuneDepot, a repository containing over 3.5 million high-quality labels (**Fig. 3A**). We hypothesized that this repository could be utilized to improve and accelerate various analyses in spatial proteomics (**Fig. 4A**).

**Figure 4.**
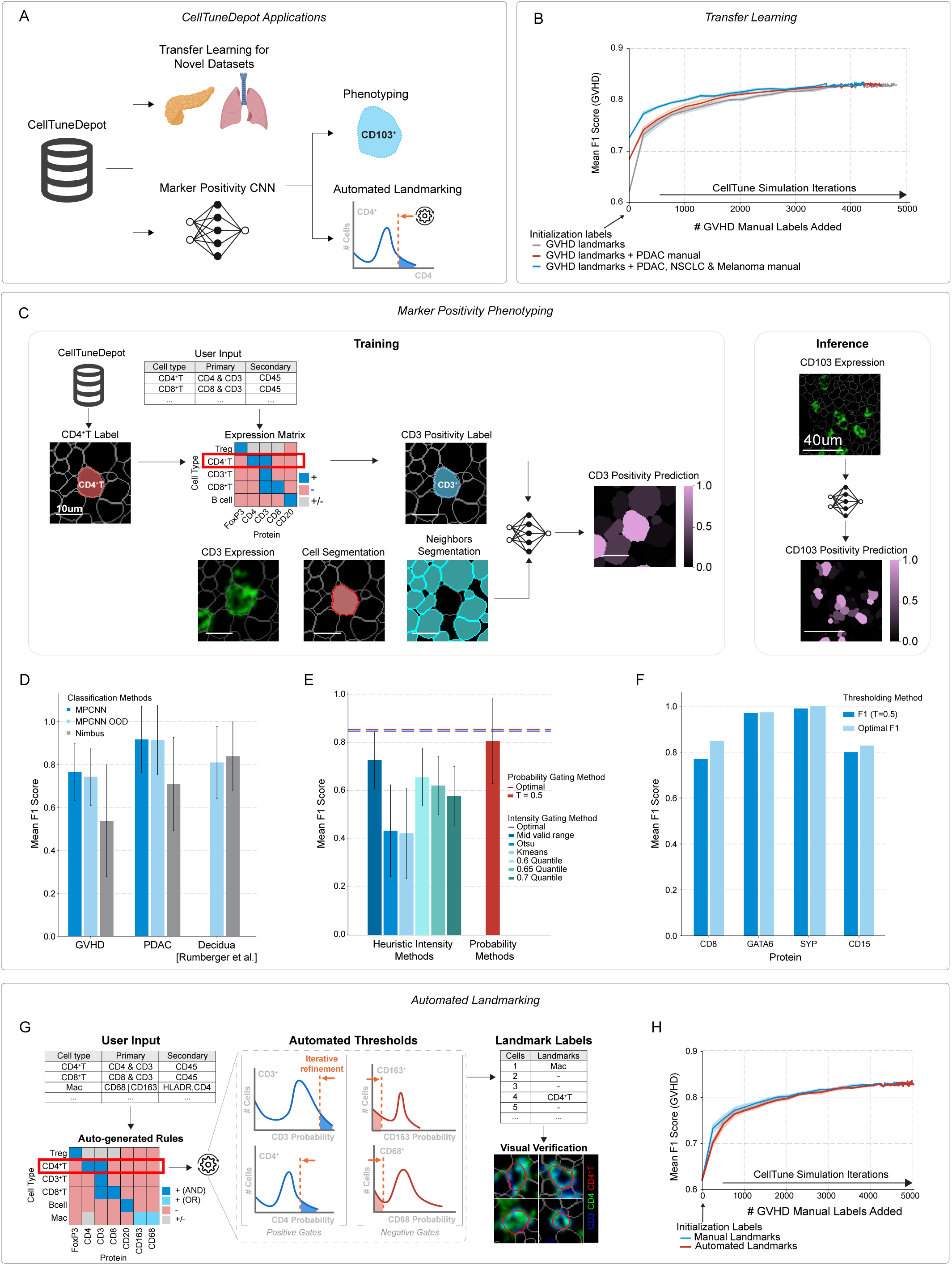
CellTuneDepot facilitates transfer learning, marker positivity prediction, and automated landmarking. **(A)** Overview schematic illustrating how CellTuneDepot provides labeled data for downstream applications, including transfer learning, phenotyping with marker positivity predictions, and automated landmark gating. **(B)** Dataset transfer: Simulation comparing model performance between models initially trained on GVHD landmark cells alone versus those trained with additional labeled data from CellTuneDepot datasets (PDAC, NSCLC, melanoma). In all cases, the initial training set includes manually gated GVHD landmark cells, followed by the incremental addition of GVHD cell labels. Model evaluation is performed on a GVHD test set, demonstrating the advantage of incorporating prior knowledge from CellTuneDepot. **(C)** Schematic of training and inference of CellTune’s marker-positivity convolutional neural network (MPCNN). To generate training data for the network, positivity labels (pos/neg) are generated automatically for different protein images. For each cell in CellTuneDepot, the corresponding cell label is used to create positivity labels for different proteins according to predefined expression associations. For example, T helper cells will be classified as CD4+, CD3+, CD20-etc. The network receives an image of one protein, centered on a cell of interest, together with segmentation masks of the cell and its neighbors. It is trained to outputs the probability of this cell to be positive for this protein. **(D)** MPCNN performance evaluation across three datasets: GVHD, PDAC, and Decidua (Rumberger et al.). For each dataset, the MPCNN is tested on: (1) Held-out images belonging to the same dataset as the training set (MPCNN, blue). 2) A novel dataset (out-of-distribution, MPCNN OOD, light blue). Its performance is compared to Nimbus (Rumberger et al.), which was trained on the Decidua dataset (gray). This data was not available for training the MPCNN. **(E)** Comparison between MPCNN-probability-based and intensity-based marker positivity classification. Optimal performance for gating on either intensity (blue dashed line) or MPCNN-probability (red dashed line) was obtained by choosing for each protein the threshold that leads to best performance on the test data. Using the default probability threshold (0.5) for MPCNN approaches optimal performance, while different heuristics to threshold the intensity fall short. **(F)** MPCNN generalizes to unseen proteins that were excluded from training, achieving high F1 scores across diverse marker types. Shown is the performance for gating the probabilities using both the default (T=0.5) and optimally-tuned thresholds. SYP = Synaptophysin. **(G)** MPCNN-probabilities can be used for automated landmarking. Users provide expected protein expression per cell type as input; the algorithm automatically generates classification rules and iteratively adjusts thresholds to identify highly confident landmark cells for each type. Landmark cells can be verified visually in the CellTune viewer. **(H)** Shown is the mean F1 score for cell classification on the GVHD dataset (y-axis). Comparison of classification accuracy (y-axis) between CellTune simulations (x-axis) initialized with manually-gated landmarks versus automated landmarks based on MPCNN probabilities demonstrates equivalent performance.

First, we investigated whether existing labeled datasets could enhance cell classification of a new dataset. We trained a model using labels from three MIBI-TOF datasets: non-small cell lung cancer (NSCLC), pancreatic ductal adenocarcinoma (PDAC), and melanoma. This model was then applied to classify the GVHD dataset, and its performance was evaluated against the manual consensus labels. To address the new set of proteins and novel cell types present in the GVHD dataset, we supplemented the labels from the three training datasets with gated landmark cells from the GVHD dataset, based on the premise that these labels are straightforward to generate. We found that the initial model that was trained exclusively on landmark GVHD labels performed worse than the model that incorporated labels from the other datasets (F1=0.62 and F1=0.73, respectively; **Fig. 4B**). To simulate the CellTune workflow, we incrementally added additional GVHD training labels and evaluated model performance at each iteration (Methods). Notably, the model pre-trained on the three external datasets achieved comparable performance to the model trained solely on GVHD labels, while requiring only approximately half as many labeled cells from the GVHD dataset (**Fig. 4B**). Overall, these results indicate that CellTuneDepot can accelerate cell classification of novel datasets.

Next, we sought to leverage the high-quality labels in CellTuneDepot to train a marker-positivity network – a model that, given an image of a protein and a cell segmentation mask, predicts the probability that the cell is positive for that protein (**Fig. 4C**). This network could assist with two tasks in spatial proteomics analysis: phenotyping and landmark gating. Phenotyping is commonly performed following cell type classification and involves determining the state of different cells in the dataset according to the expression of functional proteins. For example, cycling cells will be positive for Ki-67 and apoptotic cells will express cleaved caspase 3. While lineage-associated proteins often recur across datasets, functional proteins are more dataset-specific, making the task more challenging. Landmark gating is used to define a set of high-confidence labeled cells for initial classifier training. Both tasks require reliable marker positivity classification, which is often approximated by thresholding raw expression values but is susceptible to errors due to spillover and staining variability.

Training a generalized marker-positivity network requires positivity labels for a large set of proteins, which are cumbersome to generate manually. However, it was previously suggested ^29,40^ that given a set of labeled cells, one can easily generate automatic positivity labels for a large set of proteins, by incorporating prior knowledge on expected expression patterns. For example, a cell that is classified as a cytotoxic T cell will be classified as positive for CD8, CD3 and CD45, and negative for CD20, CD68, Keratin and more. These labels may have errors but are easy and fast to generate. We hypothesized that CellTuneDepot’s high-quality labels could enable us to train an improved marker-positivity network using this strategy. To this end, we generated a table specifying the expected positivity status for all proteins across all cell types in our datasets. We then used this table to assign protein positivity labels for every cell based on its cell type. A single convolutional neural network was trained on these labels to predict positivity for any protein, regardless of cell type or dataset. The network receives as input three images centered around a cell of interest depicting: 1. The protein stain. 2. The segmentation mask of the cell to be classified. 3. The segmentation mask of neighboring cells. The network is trained to output the positivity classification of this protein in this cell (positive/negative) according to the table (**Fig. 4C**). The datasets are greatly enriched for negative over positive labels (88% vs. 12% respectively). Therefore, this architecture, which operates at the level of a small crop centered on a cell of interest, was advantageous in enabling us to balance between positive and negative labels.

We trained the marker positivity convolutional neural network (MPCNN) on four datasets, integrated it into the software (**Fig. S2I**), and evaluated it on heldout data from the same dataset, a different dataset acquired and labeled by our lab, or a published manually-labeled dataset ^40^. We compared the results to Nimbus ^40^, a published model for marker positivity that was trained on manually labeled FOVs of just one dataset. We found that MPCNN achieved high F1 scores of 0.74-0.92 across datasets, even when evaluated on novel data. In the case of the external dataset the F1 was comparable to the F1 published by the authors using a model that was trained exclusively on the same data (**Fig. 4D**). Overall, we deduce that the MPCNN trained on diverse high-quality CellTune labels has improved performance and capability of generalization to novel datasets.

The probabilities outputted by the MPCNN correlated with, but were not identical to, protein expression (**Fig. S3A-C**). Cells with intermediate intensity levels showed the most variability in their predicted marker positivity probabilities. This variability was particularly evident in membranous markers affected by spillover (e.g., CD8; **Fig. S3D-F**), moderately observed in cytoplasmic markers (e.g., Mucin), and least pronounced in nuclear markers (e.g., FOXP3). In some cases, cells with zero measured intensity exhibited non-zero positivity probabilities due to signal detected just outside the segmentation mask. In rare instances, these probabilities exceeded the classification threshold (>0.5), suggesting that the segmentation may have excluded relevant signal (**Fig. S3G**). Conversely, cells with very high intensity may have low predicted probability attributed to spillover (**Fig. S3H**). These observations indicate that the MPCNN captures nuances beyond directly measuring intensity and may offer added discriminatory power, both for challenging intermediate-expression cells and for cells affected by imperfect segmentation.

To explore the utility of the MPCNN we compared the F1 scores of positivity classification across all proteins obtained by thresholding either the probabilities of the MPCNN or the expression values. Interestingly, we found that if we tailor the threshold of each protein to maximize the F1 score, then gating on the intensity or the positivity probability is comparable, with slightly better performance using the probability (**Fig. 4E** dashed lines). However, identifying an optimal threshold per protein is challenging and time consuming. Applying a simple threshold of 0.5 to the MPCNN probabilities yields better performance (F1=0.81, red bar in **Fig. 4E**) than any of the heuristics tested for intensity-based gating (F1=0.41-0.72, blue bars in **Fig. 4E**). Altogether, we conclude that the MPCNN trained on CellTuneDepot labels achieves good positivity classification across proteins and datasets without investing much time and effort in tailoring the threshold.

Having established the general performance of the MPCNN, we next evaluated its utility in the two key applications introduced earlier: phenotyping and landmark gating. For phenotyping, we tested whether the MPCNN could generalize to novel proteins by training it on a limited set of proteins and evaluating its performance on unseen proteins that were excluded from training. We observed high F1 scores of 0.78-1, depending on the properties of the specific protein (**Fig. 4F**), suggesting that the MPCNN could generalize to novel proteins as is needed for phenotyping.

Another prominent application of the MPCNN we explored was the automation of the landmarking step in the initialization of CellTune. To test this, we trained an MPCNN on all datasets, except for the GVHD. We then used this MPCNN to automate the process of landmark gating. Briefly, the researcher constructed a table of cell types and their expected positivity for different proteins and designated a minimum number of landmark cells for each cell type (e.g. 20 cells). The algorithm automatically generated gating schemes, and landmarked cells according to stringent thresholds on the MPCNN probabilities. The process was repeated automatically, with gradual relaxation of the thresholds until the desired number of landmark cells was reached (**Fig. 4G**, Methods). We then simulated the CellTune classification process, iteratively adding labels. We found that classification accuracy was comparable, whether initializing the model with landmarks generated by user-defined custom gates or by landmarks automatically generated from MPCNN probabilities (**Fig. 4H, Fig. S2J).** We conclude that the MPCNN trained on CellTuneDepot can accelerate landmarking of novel datasets by alleviating the need for manual gating.

Overall, we demonstrate that the high-quality labels of CellTuneDepot can be leveraged to improve classification of novel datasets, and that the MPCNN trained on these labels can substantially expedite both phenotyping and landmark gating. CellTuneDepot is released together with the CellTune software alongside a pretrained MPCNN model, enabling users to easily apply marker-positivity classification to their data. In addition, newly labeled datasets can be incorporated into CellTuneDepot to enable retraining or fine-tuning the MPCNN as needed, ensuring adaptability to evolving research needs.

### CellTune’s design and features contribute to performance

CellTune’s cell classification pipeline incorporates several innovations, including adding spatial expression features and utilizing the QBC approach to prioritize cells for human labeling. CellTune’s implementation of QBC is designed to intelligently sample cells in order to balance agreement across cell types and images while also giving extra priority to rare cell types by default, though users can customize sampling priorities if desired. We explored the effects of these features on classification performance. First, we examined whether CellTune’s QBC approach with default settings reduced the number of labels needed for accurate classification. To this end, we ran three simulations of CellTune in which the pipeline was initialized with the same landmark cells, and then additional labeled cells were added in each round (Methods). In the first simulation the labeled cells were added in an FOV-wise manner, in the second random labeled cells were added and in the third the cells were added by the QBC approach (**Fig. 5A, left**). We found that adding labels by FOVs was highly inefficient (F1 at 5000 cells = 0.78), presumably since many cells from the same FOV hold similar information, which doesn’t necessarily translate to other FOVs, resulting in model overfitting. Adding labeled cells randomly increased performance (F1 at 5000 cells = 0.85), but CellTune’s QBC performed best (F1 at 5000 cells = 0.94) (**Fig. 5A, right**). Indeed, when we examined the neighborhoods of the cells that were manually labeled using the QBC approach, we found that manually labeled cells are over-represented in neighborhoods with a high diversity of cell types (**Fig. 5B**), suggesting that the QBC approach prioritizes “difficult” cells with high information content. Furthermore, we observed that the automatic balancing within CellTune’s QBC approach resulted in cell type frequencies in manual labels that were more evenly distributed than their their representation in the dataset (**Fig. 5C**), effectively addressing the high imbalance of labels typically found in tissue images. Together, these results suggest that CellTune’s QBC sampling strategy promotes labeling of diverse and difficult cells, overall improving accuracy while minimizing the number of cells for manual labeling.

**Figure 5:**
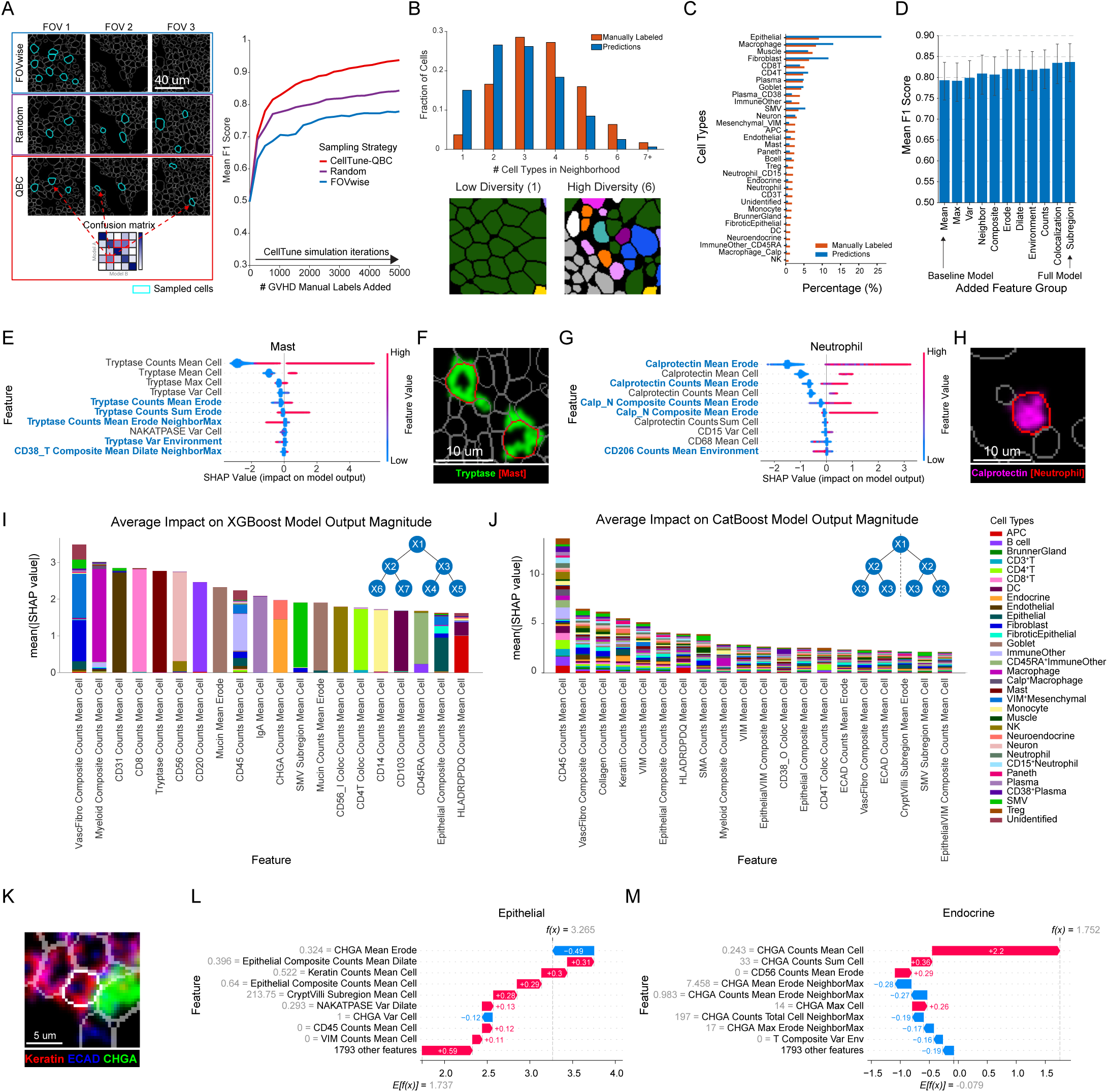
CellTune’s design and features contribute to performance. **(A)** (Left) Schematic comparing different sampling strategies for labeling. Query-by-committee (QBC), used in CellTune, is compared to random sampling and sampling by field-of-view (FOV). (Right) Simulations of iterations of model-training and labeling. Cells are selected for labeling using the different sampling strategies. Mean F1 classification scores (y-axis) are calculated as cell labels are added to the training set (x-axis). The initial training set (# added cells = 0) includes the gated landmark cells for each cell type (see Methods for simulation details). **(B)** Top: Neighborhood diversity compared between cells that were sampled for manual labeling by CellTune (orange) versus those that were not (blue). Bottom: Examples of cells in low and high diversity neighborhoods. **(C)** Histograms showing the percentage of different cell types in the dataset (Predictions, blue) and in the subset prioritized for labeling by CellTune (Manual, orange). **(D)** Mean F1 scores for models trained by sequentially adding feature groups to the baseline model in order of increasing feature complexity. Each bar represents a different model, with the baseline model (first bar) using only the mean expression, while subsequent models incorporate additional feature groups cumulatively. The final bar represents the full model incorporating all 1802 features. Error bars indicate 95% confidence intervals across all cell types. **(E)** SHAP interpretation of the XGBoost model, showing the top 10 features contributing to the prediction of Mast cells. Features are ranked by their mean absolute SHAP values, with higher values indicating greater importance. Each point represents a SHAP value for a single prediction, with color indicating the feature value (e.g., low to high). Spatial features are highlighted in blue. **(F)** Example of tryptase expression in mast cells spilling over into neighboring cells. **(G)** SHAP Top 10 features contributing to the prediction of Neutrophil cells. **(H)** Example of Calprotectin expression in the center of a neutrophil. **(I-J)** Impact of the 20 top contributing features on the model predictions for each cell type. Results shown for (I) XGBoost and (J) CatBoost. Schematics of their tree-building strategies are shown at the top right for both models. **(K)** Example of a cell misclassified by clustering as endocrine and correctly classified by CellTune as epithelial. **(L-M)** SHAP waterfall plot illustrating the XGBoost decision-making for the (L) epithelial and (M) endocrine prediction probabilities for the cell shown in (K).

Next, we examined the contribution of the spatial features to model performance by performing an ablation study. We compared the performance using the full set of CellTune’s computed features (full model) to the performance when excluding spatial information and using only the mean cell expression (Baseline). We grouped the different features into groups of related features (Methods, **Fig. S1B**), and added one feature group at a time according to the computational difficulty in generating the features, from the easiest (Max/Var) to the most difficult and/or time consuming (Subregion). We found that the full model outperformed the baseline model, highlighting the contribution of spatial features to the cell classification task (F1=0.84 vs. F1=0.79, respectively **Fig. 5D**). Moreover, we found that different features were important for different cell types (**Fig. S4**). For example, information on neighboring cells (Neighbor) was important for classification of Mast cells (**Fig. 5E, S4**), presumably since the expression of tryptase is very strong and tends to bleed into other cells (**Fig. 5F**). Similarly, the expression in the center of the cell (Erode) was important for the classification of Neutrophils (**Fig. 5G,H, S4**). Different proteins have different staining patterns, and these results suggest that CellTune learns to utilize different features in its classification to accommodate said differences.

To better understand what each model learns and how it affects the final prediction, we performed SHAP (SHapley Additive exPlanations) feature importance analysis^50^. SHAP assigns an importance value to each feature for a particular prediction. We first examined the top contributing features in the dataset, and their contribution to the prediction of each class (**Fig. 5I,J**). The two models that we used, XGBoost and CatBoost, learnt differently. In XGBoost, features primarily contributed to their associated cell types (e.g., CD8 pixel counts contributed most to CD8^+^T classification), whereas in CatBoost, every feature contributed similarly across multiple cell types. This difference arises from the architectures of the decision trees: CatBoost builds symmetric trees, forcing every prediction to use multiple features, while XGBoost builds asymmetric trees, where each prediction uses only required features. Using two different architectures ensures that the models learn differently, enhancing the utility of the QBC approach.

Examining the SHAP values for individual predictions further highlights the importance of incorporating spatial features to address challenges like signal spillover. Figure 5K shows an example of a cell that was classified as epithelial by CellTune, but as endocrine by clustering. Visual inspection shows that this cell is neighboring an endocrine cell with high expression of chromogranin, which is spilling into the cell of interest. SHAP waterfall plots clarify the decision-making process of the XGBoost model. For the epithelial prediction (**Fig. 5L**), chromogranin expression contributed negatively but were outweighed by epithelial-specific features, such as keratin. In contrast, for the endocrine prediction (**Fig. 5M**), chromogranin expression within the cell’s segmentation (Erode) positively contributed to the classification, but the chromogranin signal from neighboring cells (NeighborMax) had a stronger negative influence on the prediction. Overall, we conclude that incorporation of spatial features improves classification.

## Discussion

Accurate cell classification is essential for analyzing spatial proteomics tissue images, as it underpins downstream analyses. Misclassification of cell types can compromise results, yet this task is challenging due to signal spillover, irregular cell shapes, dense tissue packing, and the two-dimensional imaging of three-dimensional structures. Additional hurdles include imbalanced data, where cell type frequencies and marker proteins vary significantly, and technical artifacts such as tissue quality variations and imaging noise.

A robust approach to cell classification must address these challenges. Leveraging spatial expression features, including subcellular localization, protein colocalization, and neighborhood patterns, is critical to disentangle real biological signals from spillover artifacts. Prior biological knowledge is imperative for this task, yet the timing and method of its infusion can impact classification outcomes. Predefining cell types or using labeled examples can guide the model but may limit the discovery of novel cell types. Conversely, post-classification manual curation enables novel findings but is labor-intensive and does not enhance the algorithm during training.

CellTune addresses these challenges by combining several innovations into a unified framework specifically designed for cell classification in multiplexed imaging. Its strength lies in the integration and optimization of (1) a human-in-the-loop active learning workflow, (2) spatial feature extraction capturing subcellular and neighborhood-level information, (3) intelligent cell sampling with query-by-committee, and (4) an interface for visualization, annotation, gating, and classification at scale. This synergistic design yields both higher classification accuracy and enhanced cell type resolution compared to alternative methods; and achieves both with limited manual effort, even for datasets containing hundreds of images and millions of cells. Importantly, CellTune balances the incorporation of prior biological knowledge with the ability to uncover new cell types. By integrating spatial features into the classification process and prioritizing uncertain or conflicting cases for manual labeling, CellTune minimizes labeling effort while improving accuracy. This iterative approach, inspired by active learning tools for low-plex imaging^51,52^ and query-by-committee strategies from other domains^53–56^, focuses labeling efforts on the most informative cells, enhancing model performance and enabling finer-grained classification.

CellTune was designed and refined through direct application to large, diverse datasets, in close collaboration with end-users, ensuring that the system meets the practical needs of researchers working with multiplexed imaging data. It is well-suited for real-life data analysis scenarios, such as revisiting cell classification due to newly added samples, missed cell types, or overlooked artifacts. Its modular design allows models trained on one dataset to be adapted to others, even when marker overlap is partial, reducing the need for retraining and manual labeling. CellTune is robust enough to perform well with default settings, while remaining flexible: users can easily introduce new cell types at any stage and incorporate custom features, whether tabular or image-derived.

Although no architectural innovations were made to the underlying classifiers, XGBoost and CatBoost, their state-of-the-art performance within CellTune arises from their combination with spatial features and high-quality labels generated through iterative active learning. By focusing manual annotation on the most informative cells, CellTune produces training data that maximizes classifier effectiveness. While manual annotation remains time-consuming, CellTune’s optimized graphical interface significantly streamlines the process, making expression data easily accessible and facilitating cell population management tasks. Typical projects with hundreds of images and over a million cells required ±80 hours, yielding high-quality labels that outperformed ten state-of-the-art methods. The entire workflow is designed for practical usability, where the time investment scales with the diversity and complexity of cell types rather than the sheer number of cells. As more labeled datasets are generated, annotation time for new projects is expected to decrease further.

While several existing tools such as QuPath^37^, napari^38^, MCMICRO^39^, ilastik^51^, and Mantis^57^ support aspects of multiplexed imaging analysis, none provide a fully integrated solution tailored to rapid and accurate cell classification at scale. QuPath, while widely used for histopathology, requires users to manually train classifiers for each marker and lacks optimized support for large, multiplexed cohorts. Napari, although versatile, is a general-purpose viewer not specifically designed for multiplexed imaging, and often struggles with stability, scalability, and usability in this context. MCMICRO offers an extensive pipeline for preprocessing but lacks an integrated interface for efficient cell-level annotation and active learning, relying on modules composed of external tools like scimap^58^ for clustering and gating. Mantis provides a promising interface, but it also struggles with stability and is severely limited in its capacity to run classification algorithms or compute spatial features at scale. Ilastik includes an active learning framework and limited spatial features, but it lacks the capacity to handle multiplexed images effectively, and its performance degrades significantly with increasing dataset sizes. In contrast, CellTune integrates active learning, spatial feature computation, and advanced visualization into a single, scalable software package. Its graphical interface is designed explicitly for efficient labeling of multiplexed images, with intelligent channel management, population tracking, and seamless transition between labeling, model training, and result inspection. While the core components of the active learning framework could, in principle, be adapted into other platforms like napari or QuPath, CellTune’s implementation is optimized specifically for the challenges of large, high-dimensional multiplexed imaging datasets and offers significant practical advantages out-of-the-box.

Beyond the software and method itself, a key contribution of this work is CellTuneDepot, a large repository comprising over 40,000 manually labeled cells, including 10,000 cells annotated by three independent experts, along with an additional 3.5 million high-confidence CellTune-generated labels across six datasets. Previous ground truth efforts often focused on “easy” cells or small image subsets, limiting real-world applicability. In contrast, CellTuneDepot was constructed by randomly selecting image crops and systematically labeling all cells within them, ensuring a representative and comprehensive dataset. To address labeling fatigue and human error and variability, consensus labels from three annotators were used for a subset of cells, and comparisons with CellTune’s predictions demonstrated its ability to guide users effectively, reducing errors and increasing accuracy.

To support marker positivity training and evaluation, we generated 16.5 million marker-positivity labels from 2.7 million cells, using CellTuneDepot’s high-confidence labels, which are equivalent in quality to the manually curated “gold-standard” labels used by other methods. For example, Nimbus generated approximately 1 million manually curated marker-positivity labels, comparable in quality but considerably smaller in scale. Other attempts to construct larger datasets have relied on labels inherited from previous studies, which were predominantly generated by clustering-based methods with only limited manual curation, if any. In our benchmarking against our dataset annotated by three experts, which we consider as our gold-standard, we found that such clustering-based labels yielded poor classification performance without extensive correction — even under an optimal scenario where we assigned the best possible label to each cluster. In contrast, CellTune and CellTuneDepot uniquely combine scale with high-quality label generation, providing a valuable resource for robust and generalizable model training and evaluation.

CellTune and CellTuneDepot together offer a powerful framework for cell classification, integrating visualization, analysis, and data resources. Built as a modular Python-based platform, CellTune is easily expandable, allowing for the integration of new algorithms, classifiers, and features as the field evolves. This adaptability guarantees it remains a dynamic tool, pushing the boundaries of automated cell classification while ensuring flexibility and reliability. With its long-term vision of fully automated, biologically informed classification, CellTune and its extensive labeled datasets provide an invaluable resource for advancing spatial proteomics and tissue analysis.

## Methods

### Datasets, cell types, and labels

#### Graft-versus-host disease (GVHD) MIBI dataset

The GVHD MIBI dataset consists of 75 0.4×0.4 mm^2^ images taken from ref.^59^. The 75 fields of view (FOVs) include tissues of 13 GVHD patients and 10 controls, altogether encompassing a total of 159,516 cells. Cells were classified with CellTune into the following 32 cell types (primary marker expression definition in parentheses): Treg (FOXP3^+^), CD8^+^T (CD8^+^), CD4^+^T (CD3^+^CD4^+^), CD3^+^T (CD3^+^), B cell (CD20^+^), Plasma (IgA^+^), CD38^+^Plasma (CD38^+^IgA^-^), Mast (Tryptase^+^), Neutrophil (Calprotectin^+^), CD15^+^Neutrophil (CD15^+^Calprotectin^-^CD68^-^CD163^-^CD206^-^), Calprotectin^+^Macrophage (Calprotectin^+^(CD68^+^|CD163^+^| CD206^+^)), Macrophage (CD68^+^|CD163^+^|CD206^+^), Dendritic Cell (CD103^+^CD45^+^HLADRDPDQ^+^), Antigen Presenting Cell (CD45^+^HLADRDPDQ^+^), Monocyte (CD14^+^), ImmuneOther (CD45^+^), CD45RA^+^ImmuneOther (CD45RA^+^), Goblet (Mucin^+^), Paneth (Lysozyme^+^), Brunner Gland (Lysozyme^+^ in BrunnerGland Subregion), Fibroblast (SMA^+^|Collagen^+^), Fibrotic Epithelial (ECAD^+^(SMA^+^|Collagen^+^)), Smooth Muscle Vasculature (SMA^+^ in Vasculature Subregion), Muscle (SMA^+^ in Muscle Subregion), Endothelial (CD31^+^), Neuron (CD56^+^CD45^-^), NK (CD56^+^CD45^+^), Epithelial (ECAD^+^|Keratin^+^|NaKATPASE^hi^), Endocrine (CHGA^+^), Neuroendocrine (CD56^+^CHGA^+^), Mesenchymal (VIM^+^), and Unidentified (negative for all lineage markers).

48 crops were selected from different FOVs of the GVHD MIBI dataset for manual labeling to generate a ground-truth for testing. Crops were of size 120×120 μm^2^, each encompassing ∼200 cells, for a total of 10,283 cells. The first 15 crops were randomly selected (both FOV and location), while the remaining 15 were chosen to enrich rare cell types (based on preliminary gating). To achieve this, we repeatedly sampled random locations from the remaining FOVs until a crop contained predictions for at least one of the rarest cell types. Two labelers were tasked with manually labeling all the cells in the crop from scratch. A third (most expert) manual labeler was tasked with labeling the cells that the two labelers disagreed on, as well as with reviewing, curating, and establishing a consensus for all the labels. Labelers struggled with labeling all 32 cell types classified with CellTune within a reasonable timeframe, thus we reduced the resolution from 32 to 18 cell types for manual labeling and tested by combining multiple cell types as follows (the resulting combined primary marker expression rule is shown in parentheses): Monocyte was combined with Macrophage (CD14^+^|CD68^+^|CD163^+^|CD206^+^), CD38^+^Plasma with Plasma (IgA^+^|CD38^+^), Neuroendocrine with Endocrine (CHGA^+^), Brunner Gland with Paneth (Lysozyme^+^), Mesenchymal with Unidentified (VIM^+^|negative for all lineage markers), NK, Dendritic Cell, Antigen Presenting Cell, and CD45RA^+^ImmuneOther were combined with ImmuneOther (CD45^+^|CD45RA^+^), Smooth Muscle Vasculature, Muscle, and Fibrotic Epithelial were combined with Fibroblast (SMA^+^|Collagen^+^), and CD15^+^Neutrophil and Calprotectin^+^Macrophage were combined with Neutrophil (CD15^+^|Calprotectin^+^).

#### Colorectal cancer (CRC) CODEX dataset

Imaging data for the CRC CODEX dataset were taken from ref.^60^. From their TMA, we selected one of the 0.6-mm cores per patient, yielding a total of 35 FOVs. We used Mesmer^33^ to perform segmentation, which resulted in a total of 94,389 cells. Cells were classified with CellTune into the following 17 cell types (primary marker expression definition in parentheses): Treg (FOXP3^+^), CD8^+^T (CD8^+^), CD4^+^T (CD3^+^CD4^+^), CD3^+^T (CD3^+^), B cell (CD20^+^), Plasma (CD38^+^), Neutrophil (CD15^+^), Macrophage (CD68^+^|CD163^+^|CD11b^+^), Dendritic Cell (CD11c^+^), Fibroblast (SMA^+^), Smooth Muscle Vasculature (SMA^+^ in Vasculature Subregion), Endothelial (CD31^+^), Neuron (SYP^+^|CD56^+^), Lymphatic (PDPN^+^), Tumor (Cytokeratin^+^|NaKATPASE^+^| MUC1^+^), Stroma (VIM^+^), and Unidentified (negative for all lineage markers).

Eight crops were selected from different FOVs of the CRC CODEX dataset for manual labeling to create a ground-truth for testing. Each crop measured 200×200 μm^2^ and contained ∼350 cells, totaling 2,745 cells. Initially, the eight crops were randomly selected, followed by three FOV swaps and adjustments to crop locations to ensure adequate representation of all cell types (based on preliminary gating). One labeler assigned labels to all cells from scratch using the same 17 cell types described above, while a second labeler reviewed and curated the annotations.

#### Breast cancer CODEX dataset

Imaging data and segmentations for the Breast CODEX dataset were taken from ref.^61^. From their test set, we selected 30 TMA cores (1.0-mm diameter each) and randomly extracted a 200×200 μm^2^ crop from each for manual labeling. Each crop contained ∼250 cells, totaling 7,610 cells overall. One labeler assigned labels to all cells from scratch using the following 14 cell types (primary marker expression definition in parentheses): CD8^+^T (CD8^+^), CD4^+^T (CD3^+^CD4^+^), CD3^+^T (CD3^+^), B cell (CD20^+^), Plasma (CD38^+^), Neutrophil (CD15^+^), Mast (Tryptase^+^), Antigen Presenting Cell (HLADRDPDQ^+^), Macrophage (CD68^+^), Monocyte (CD14^+^), Fibroblast (SMA^+^|COL1A1^+^|FAP^+^), Endothelial (CD31^+^), Tumor/Epithelial (Pan-keratin^+^|ECAD^+^|Cytokeratin7^+^|Cytokeratin5^+^), and Unidentified (negative for all lineage markers). A second labeler reviewed and curated the annotations.

#### Melanoma MIBI dataset

The melanoma MIBI dataset is an unpublished dataset consisting of 302 FOVs (0.5×0.5 mm²) from metastatic melanoma tissues of 50 patients across various anatomical sites, including lung, liver, lymph node, and skin, as well as three control tissues (melanoma, lymphoma, and kidney). In total, the dataset contains 577,798 cells. Cells were classified with CellTune into the following 20 cell types, along with one additional “Garbage” class for cells expressing multiple conflicting lineage markers due to staining artifacts missed in low-level processing (approximately 1 in 2,000 cells): B cell (CD20^+^), CD4^+^T (CD3^+^CD4^+^), CD8^+^T (CD8^+^), CD3^+^T (CD3^+^), Treg (FOXP3^+^), NKT (CD3^+^(CD56^+^|CD57^+^)), NK (CD3^-^(CD56^+^|CD57^+^)), Neuron (CD56^+^CD45^-^), Neutrophil (MPO/Calprotectin^+^), Hemosiderin Macrophage (iNOS^+^CD20^+^Fe56^hi^), Myeloid (CD16^+^|CD68^+^|CD163^+^|CD206^+^|CD209^+^|CD11c^+^|iNOS^+^|(HLADRDPDQ^+^(CD45^+^|CD45RO^+^|CD14^+^))), ImmuneOther (CD45^+^|CD45RO^+^|CD14^+^), Activated Fibroblast (FAP^+^), Myofibroblast (SMA^+^), Smooth Muscle Vasculature (SMA^+^ in Vasculature Subregion), Endothelial (CD31^+^), Epithelial (BCAT^+^ in Epithelial Subregion), Tumor (SOX10^+^), BCAT^+^Tumor (BCAT^+^SOX10^-^), Unidentified (negative for all lineage markers), and Garbage (artifact staining).

Ten FOVs were selected for manual labeling to create a ground truth for testing. Each entire FOV (∼2,000 cells per FOV) was labeled by a single manual labeler with the same cell type definitions described above, totaling 21,204 labeled cells. FOVs were chosen by an expert to ensure a wide range of staining quality, tissue types, and cell types based on preliminary gating.

#### Non-small-cell lung cancer (NSCLC) MIBI dataset

The NSCLC MIBI dataset is an unpublished dataset consisting of 302 FOVs from a cohort of 20 early-stage NSCLC patients (stage I–II) who were treated with neoadjuvant pembrolizumab (anti-PD-1) as part of a Phase I clinical trial. Core-needle biopsies (CNBs) and tumor resections were imaged as part of the study. The dataset includes: 161 FOVs at 0.2×0.2 mm², 60 FOVs at 0.4×0.4 mm², and 81 FOVs at 0.8×0.8 mm². In total, the dataset contains 706,550 cells, classified with CellTune into 30 cell types and one additional Garbage class for artifacts resulting from staining errors (approximately 1 in 312 cells). The classified cell types and their primary marker definitions are as follows: Treg (FOXP3⁺), CD8⁺T (CD8⁺), CD4⁺T (CD4⁺CD3⁺), CD3⁺T (CD3⁺), CD8⁺Treg (FOXP3⁺CD8⁺), NKT (CD56⁺CD3⁺), NK (CD56⁺CD45⁺), B cell (CD20⁺), Germinal B (CD21⁺CD20⁺), IgA⁺Plasma (CD38⁺IgA⁺), Plasma (CD38⁺IgA⁻), CD31⁺Plasma (CD31⁺CD38⁺), Mast (Tryptase⁺), Calp⁺Macrophage (Calprotectin⁺(CD14⁺|CD68⁺|CD163⁺|CD206⁺)), Neutrophil (Calprotectin⁺), Macrophage (CD68⁺|CD163⁺|CD206⁺), Dendritic Cell (DC_LAMP⁺|CD103⁺HLADRDPDQ⁺| CD11c⁺), Monocyte (CD14⁺), ImmuneOther (CD45⁺), CD103⁺ImmuneOther (CD103⁺), Antigen Presenting Cell (HLADRDPDQ⁺), Pneumocyte Type 2 (DC_LAMP⁺(EpCAM⁺|Keratin⁺)), Smooth Muscle Vasculature (SMA⁺ in Vasculature Subregion), Fibroblast (SMA⁺|Collagen⁺), Endothelial (CD31⁺), Neuron (CD56⁺CD45⁻), Epithelial (EpCAM⁺|Keratin⁺), Mesenchymal (VIM⁺), IgA⁺Unidentified (IgA⁺CD20⁻CD38⁻), Unidentified (negative for all lineage markers), and Garbage (artifact staining). CD103⁺ImmuneOther was subsequently merged with ImmuneOther due to very low cell counts. The remaining 29 cell types are shown in Figure 3I and their mean expression is shown in Figure 3J.

#### Pancreatic ductal adenocarcinoma (PDAC) MIBI dataset

The PDAC MIBI dataset is an unpublished dataset consisting of 618 FOVs (0.4×0.4 mm²) from 117 patients with pancreatic ductal adenocarcinoma (PDAC). In total, the dataset contains 1,252,322 cells, classified with CellTune into 27 cell types – cell types and their primary marker definitions are as follows: Treg (FOXP3⁺), CD8⁺T (CD8⁺), CD4⁺T (CD4⁺CD3⁺), CD3⁺T (CD3⁺), B cell (CD20⁺), Neutrophil (CD15⁺), Calprotectin⁺Neutrophil (Calprotectin⁺), Antigen Presenting Cell (HLADRDPDQ⁺CD45⁺), Monocyte (CD14⁺), ImmuneOther (CD45⁺), Fibroblast (SMA⁺|COL1A1⁺|FAP⁺), GATA6⁺Fibroblast (GATA6⁺(SMA⁺|COL1A1⁺|FAP⁺)), Smooth Muscle Vasculature (SMA⁺ in Vasculature Subregion), Endothelial (CD31⁺), Neuron (SYP⁺VIM⁺), Endocrine Alpha (SYP⁺FAP⁺), Endocrine Other (SYP⁺), Acinar (Amylase⁺), Classical Tumor (Keratin⁺GATA6⁺), Basal Tumor (KRT5⁺), Mixed Tumor (GATA6⁺KRT5⁺), Tumor (Keratin⁺), Mesenchymal (VIM⁺), GATA6⁺Mesenchymal (VIM⁺GATA6⁺), Muscle (SMA⁺ in Muscle Subregion), Epithelial (ECAD⁺Keratin⁻Amylase⁻SYP⁻), and Unidentified (negative for all lineage markers).

#### Melanoma brain metastases (MBM) MIBI dataset

The MBM MIBI dataset is an unpublished dataset consisting of 202 FOVs (0.8×0.8 mm²) from patients with brain metastatic melanoma, including both cranial and extracranial lesions (e.g., skin, lymph node, stomach). In total, the dataset contains 999,400 cells, classified with CellTune into 29 cell types – the classified cell types and their primary marker definitions are as follows: Treg (FOXP3⁺), CD8⁺Treg (FOXP3⁺CD8⁺), CD4⁺T (CD4⁺CD3⁺), CD8⁺T (CD8⁺), CD3⁺T (CD3⁺), B cell (CD20⁺), Neutrophil (Calprotectin⁺), Macrophage (CD68⁺|CD163⁺|CD206⁺), Dendritic Cell (CD11c⁺), Antigen Presenting Cell (HLADRDPDQ⁺), Monocyte (CD14⁺), ImmuneOther (CD45⁺), Fibroblast (SMA⁺|Collagen⁺), Endothelial (CD31⁺), Smooth Muscle Vasculature (SMA⁺ in Vasculature Subregion), Neuron (CD56⁺CD45⁻), Astrocyte (GFAP⁺), Activated Astrocyte (GFAP⁺HLADRDPDQ⁺), GFAP⁺Unidentified (GFAP⁺ not in Brain Subregion), Tumor (SOX10⁺|MelanA⁺), CD56⁺Tumor (CD56⁺(SOX10⁺|MelanA⁺)), BCAT⁺Tumor (BCAT⁺ in Tumor Subregion), BCAT⁺Unidentified (BCAT⁺ not in Tumor Subregion), NK (CD56⁺CD45⁺), NKT (CD56⁺CD3⁺), Epithelial (Keratin⁺), Microglia (IBA1⁺), IBA1⁺Unidentified (IBA1⁺ not Brain Subregion), and Unidentified (negative for all lineage markers).

High-quality labels generated using CellTune on published datasets, along with all fully manually labeled crops, are available in the CellTuneDepot repository (see Data Availability). Unpublished datasets will be added to CellTuneDepot upon publication of their respective manuscripts.

### Cell classification with CellTune

The complete cell classification workflow from raw protein images to final cell type classifications is depicted in Figure S1A. CellTune does not yet include low-level processing and segmentation stages. For mass-based imaging data (MIBI), low-level processing was performed with MAUI^62^. For fluorescence imaging data (CODEX), images were cleaned using Ilastik’s pixel classification module^51^. Segmentation for all datasets was performed with Mesmer^33^ (v0.11 and later).

#### Compute features

The first step in most cell classification algorithms is feature extraction, in which the mean expression of each protein in each cell is computed. CellTune begins with a similar feature extraction step, but it computes many more features, to account for various spatial expression patterns that can aid the computational models in cell classification **(Fig. S1B)**. Each feature is derived from a channel, including single protein expression, multiple protein colocalization, composites of proteins, and subregions (via pixel classification or manual drawing). Colocalizations are computed by masking a channel by other channels. For example, a CD3-CD4 colocalization feature image masks the CD3 channel in all the pixels where CD4 is positive. This colocalization feature helps us to classify CD4^+^T cells, where we expect both CD3 and CD4 to be expressed in many of the same pixels. (Note this requires that the images were cleaned in the low-level processing, otherwise all pixels will be “on” due to background.) Composites of proteins are computed by taking the maximum value at each pixel for a set of proteins. For example, a ‘macrophage’ composite feature image can be calculated from CD68, CD163, and CD206. Larger-scale anatomical features called subregions add contextual histological information that can help distinguish between different cell types. For example, a pixel classifier which identifies smooth muscle around vessels can help distinguish between fibroblasts and smooth muscle vasculature cells^63^, which both express SMA. Currently, we use Ilastik^51^ to train pixel classifiers and generate these masks. Four statistical measures (mean, maximum, variance, and pixel count) are computed across distinct cellular compartments: the cell center (erode), entire cell segmentation (cell), immediate surroundings (dilate), and broader cellular environment (environment). By default, erode is a 5-pixel erosion (∼2 μm), dilate is a 10-pixel dilation (∼4 μm), and environment is a 50-pixel dilation (∼20 μm) of the cell segmentation **(Fig 1B left)**. These default values worked well for all our images (20X magnification, 0.4-0.5 μm per pixel) but could easily be adjusted in advanced settings if needed. Optionally probability features are computed using the MPCNN, but only on the entire cell segmentation (cell). The maximum feature values among each cell’s neighbors are additionally computed. In total, approximately 2000 features are computed per cell, depending on the number of channels, which is project-specific, then merged and stored in a large cell table **(Fig. 1B, S1B)**. CellTune’s modularity readily accommodates the incorporation of new types of features, should these be necessary.

#### Initial data examination & knowledge integration

To train an initial classification model, it is necessary to identify the expected cell types in the data and initialize some labels for each class. Experimentalists often have well-informed expectations and prior knowledge about their data, which can be further validated through unsupervised clustering. In CellTune, cells are clustered^22^ based on their mean intensity features, and clusters are examined by their median protein intensities to identify any unexpected cell types that may already be apparent (**Fig. S1C, S2A-C**). Users integrate their initial data observations and prior knowledge to compile a table listing all cell types along with their defining lineage markers: *primary markers* (proteins that must be expressed and determine cell identity) and *secondary markers* (proteins whose presence or absence does not influence classification). For example, CD3 and CD4 must both be expressed on CD4^+^T cells and are listed as primary markers, since CD3^-^CD4^+^ cells could be myeloid cells (e.g., macrophages). CD45 expression is expected on CD4^+^T cells, and all immune cells, but it is listed as a secondary marker since a CD3^+^CD4^+^CD45^-^ cell in the data should still be classified as a CD4^+^T cell. This table gets updated throughout the CellTune active learning process as cell type definitions are refined or new cell types are discovered, and it is useful for landmarking and marker positivity training data generation, as well as for standardizing cell types across projects and sharing data with collaborators and the community.

#### Manual landmarking

Unsupervised clustering generates many erroneous labels, and thus serves as a poor starting point for training. To generate high-quality labels fast, CellTune performs gating of high-confidence landmark cells **(Fig. 1B, bottom)**. It uses strict thresholds on the features derived in the previous step, both positive and negative expression, to identify landmark cells with no ambiguity. A detailed example for manually landmarking Mast cells in the GVHD dataset is shown in Figure S1D. Positive thresholds are set on the probability (and/or intensity) feature of the primary marker associated with the cell type, in this example Tryptase. In addition, a positive threshold is set on the ratio of the cell’s feature to that of its maximum neighbor, to rule out spillover with high confidence. Negative gates are set on the on the probability (and/or intensity) features of the primary markers associated with other cell types. CellTune facilitates the process of landmark gating by easily visualizing and navigating to gated cells for visual inspection and verification. Users can freely adjust the manual landmark gating thresholds, as well as add or remove any gates according to their specific data. Alternatively, the user can input initial labels generated by any other method, like by manual labeling or by using CellTune’s automated landmarking method described below.

#### Automated landmarking

We developed an automated landmarking method that automates and speeds up the landmark gating process while achieving comparable classification results. The input to the automated landmarking process is a user-generated table, that defines for each cell type: (1) a set of primary markers that define the cell type and must be present (or missing) from the cell, and (2) a set of secondary markers that might appear for this cell type. All other protein markers are assumed to be negative for the given cell type (**Fig. 4G**). The primary and secondary marker inputs allow for simple logical expressions involving multiple markers (e.g. CD4^+^T cells require both CD3 and CD4). The algorithm then converts this table into a set of marker-specific rules for each cell type. Each marker is defined as either positive, negative, or neutral.

The negative class is derived from markers that appear on the primary markers list as negative. Additionally, it includes markers that are absent from the input for a given cell type (and thus assumed negative) but appear as positive in another cell type. Using the generated per-marker rule table, a series of iteratively adjusted thresholds is used to generate lists of candidate landmark cells for each cell type. These thresholds start out very strict (e.g., >0.95 probability for positive) and are gradually relaxed. The final landmark cells are selected based on threshold levels that yield the minimum required number of cells (e.g., 20 cells per cell type).

Note that this process can utilize either the intensity levels of each marker (converted to percentiles), the probability of marker positivity generated by the MPCNN, or both (**Fig. S2J**). The suggested automated landmark cells benefit from visual verification, especially for difficult and rare cell types. Overall, this process provides a speed up over the manual process while maintaining the original performance, as shown in Figure 4H.

#### Active learning cycles

Following initialization, CellTune continues to rounds of active learning, using a human-in-the-loop and query-by-committee approach **(Fig. 1C, S1E)**. The goal of these cycles is to improve the two models’ performance, while focusing manual labeling efforts on difficult cells with high information content. Each cycle consists of the following stages: model training & inference, sampling cells, and manual annotation. During labeling, users can introduce new cell types and define additional features. Notably, this interactive process can uncover upstream pipeline issues, such as errors in low-level processing or segmentation, allowing for early troubleshooting before extensive downstream analysis. The user can continue with these cycles until they are satisfied. In our experience, users were satisfied with the classifications after labeling an average of ∼100-200 cells per cell type. At the end, the final classifications are determined by averaging the probabilities of the two models.

#### Model training, inference, and uncertainty

CellTune trains two gradient boosted decision tree models (XGBoost^44^ and CatBoost^45^) using the labeled cells (landmarks and manual labels) and the computed features in the cell table (**Fig. 1C, S1E, S2D-F**). These architectures were selected for their speed (training in 1-5 minutes on a single GPU with 10k labels, <1 min for inference), inherent incorporation of feature selection that mitigates overfitting, explainability (providing class probabilities and feature importance), and most importantly for their superior performance compared to deep learning, specifically when trained on tabular data with <100k labels^64,65^. Even so, the CellTune classification framework can easily support the incorporation of alternative models. Uncertainty is assessed by evaluating the agreement between the two models’ predictions, summarized overall, per cell type, and per field-of-view (FOV), generating a confusion matrix highlighting pairwise disagreements across the dataset. Cells with conflicting predictions are strategically sampled using the query-by-committee^43^ approach as detailed in the following section.

#### Sampling strategy

In CellTune, we aim to minimize manual labeling and direct it to cells that will have high impact on the model’s performance **(Fig. 1C)**. Intuitively, labeling cells of the same confusion has diminishing returns. If the user labeled 20 cells that were confused between ‘CD8^+^T’ and ‘B cell’, we anticipate that labeling cell 21 of the same confusion may not add a lot of information to the model, whereas instead labeling a cell confused between ‘Epithelial’ and ‘Endothelial’ will be more influential on model performance. On the other hand, adding just one label for each confusion may not suffice and users may care about certain confusions more than they care about others, depending on the specific study. With this in mind, we aimed to develop an optimal labeling strategy that can also be tailored to the specifications of each user and dataset. By default, CellTune aims to balance cell type and FOV agreement, prioritizing rare cell types, and allowing cells with user-defined specific confusions to be sampled for labeling. The user can adjust how many cells, cell types and FOVs to sample at each cycle in the process (**Fig. S2G**). For example, the user can set 100 cells with disagreements to be randomly sampled from the 5 FOVs with the lowest agreement at each cycle. In addition, the user can define that 100 cells which are confused between B cells and cytotoxic T cells will be randomly sampled across all FOVs. By default, 256 cells are sampled, including 14 cells from each of the 6 FOVs with the lowest agreement, 16 cells from each of the 7 cell types with the lowest agreement, and 10 cells sampled from each of the 6 rarest cell types (**Fig. S2H**).

Sampled cells are sorted by their predicted cell types and presented to the user sequentially, with automated channel adjustments, sorted protein expression displayed on hover, and prediction options appearing as interactive buttons for quick selection. This streamlined approach enables highly efficient manual labeling, as detailed in the Results and shown in Figure 2D.

### Phenotype Classification (Protein Positivity)

#### Marker Positivity Network (MPCNN)

CellTune’s generated high-quality labels (CellTuneDepot) enabled us to create a marker positivity convolutional neural network (MPCNN), which facilitates easy classification of marker positivity for each protein marker and for each cell (**Fig. S1A, S2I**). To train the MPCNN, we generated a dataset containing marker positivity labels. We used CellTune classifications from 4 datasets: GVHD, PDAC, NSCLC and Melanoma, consisting of a total of 1,322 FOVs. Each FOV contains 40 different marker channels, for a total of 52,880 single channel intensity images. The total number of cells across all images is 2,740,325 cells.

The positivity labels were generated automatically and were derived from cell type classifications. For most cell types, a definite marker status (positive/negative) can be assumed for a subset of the markers. For example, a cell classified as a CD4^+^T by CellTune was given CD3 positive and CD4 positive labels, and a CD8 negative label. The total number of labels generated in this manner is smaller than the theoretical maximum (i.e., number of cells × number of channels), summing up to 16,509,038 individual positivity labels. This dataset is highly unbalanced, as most cells are negative for most markers, with only 12% of the labels being positive. A weighted random sampler was used to balance the dataset during training, ensuring a more even distribution of positive and negative samples. Moreover, many of the negative examples are trivial, since they do not contain any signal. Therefore, the weights were constructed to enrich for difficult cases and ensure that at least 20% of the negative samples contained some signal. The dataset was split into a 75% training set and a 25% test set. Full FOVs were assigned to either the training set or the test set such that all FOVs from the same patient were kept together in either train or test. Separating by patients prevents data leakage and constructs a more realistic test set.

The input to the network represented a specific marker in a single cell. The input structure consisted of three channels: (1) the intensity image of the query protein, (2) the segmentation mask of the cell of interest, and (3) the segmentation mask for all the other cells in the crop (**Fig. 4C**). All these input images were created by taking a center crop of size 128×128 pixels around the cell of interest. The 3-channel input image was scaled by a factor of 32 before entering the network. The network architecture is based on a ResNet18 CNN, with an additional fully connected output layer that produces a classification result. It was trained using Adam optimizer (lr=4e-4), a BCE loss and an exponentially decaying learning rate scheduler (gamma=0.9). The network is trained for 10 epochs, with each epoch loading 250,000 cells selected from the training data using weighted random sampling. On a single V100 GPU, training took approximately 12 hours. MPCNN was implemented using PyTorch^66^ (2.6). The network’s performance was evaluated by calculating F1 scores for each of the protein markers across cells.

### Benchmarking Cell Classification Methods

Existing cell classification methods fall along a spectrum between supervised and unsupervised approaches. To benchmark both types, we divided our 10k gold-standard GVHD labels into approximately equal training and test sets. The FOVs were initially split randomly to either train or test, followed by selective reassignment to balance cell type representation and total cell counts across both sets. This design ensures that supervised approaches are trained on curated, high-quality manual labels from the same dataset, encompassing all cell types. Importantly, this does not reflect real-world applications, where training labels typically originate from different datasets, consist of unrepresentative cells that are easy to gate, or contain noise from clustering-based annotations. Consequently, our benchmarking represents an upper bound on the methods’ performance.

Similarly, to enable a fair comparison, we sought to establish an upper bound on performance for semi-supervised and unsupervised approaches. For each cell classification method we benchmarked, we tested multiple parameter configurations, ran multiple trials, and retained the highest-scoring result. Except for pixel clustering & curation, we first evaluated all methods using their default parameters and then refined their settings over 5–10 hours to optimize performance. (This excludes training runtime and the initial time required for installation and testing on example datasets, where applicable.) For steps requiring threshold selection (e.g., for gating or clustering), we used the ground truth test labels to determine the optimal threshold that maximized performance. In real-world applications, users typically rely on heuristics and cannot perfectly tune thresholds, meaning our approach again represents an upper bound on the methods’ performance. Overall, we are confident that our benchmarking experiments provided a fair and rigorous evaluation of all methods. Further details and the final parameters used for each method are provided below.

#### MAPS

MAPS^31^ is a supervised feed-forward neural network for cell classification. Of all methods, MAPS was the easiest to set up, test, optimize, and run in parallel, requiring the least effort for parameter tuning. It was trained on the GVHD training set. Multiple hyperparameter configurations were tested, including models trained on only lineage markers vs. all markers + area. The final model achieving the best performance on the test set was trained on all markers + area with batch size = 128, max epochs = 500, min epochs = 250, and patience = 100.

#### STELLAR

STELLAR^27^ is a geometric deep learning framework that integrates molecular and spatial features to classify known cell types and discover novel ones. It constructs a spatial graph based on cell proximity and applies a graph convolutional neural network to learn cell embeddings for annotation. The model constructs a graph representation where each cell is embedded based on its molecular features and spatial relationships. By incorporating unlabeled cells, STELLAR can learn a joint embedding that captures both spatial and molecular similarities between cells, enabling it to classify known cell types while also discovering novel cell types in the embedding that do not fit into the predefined categories. We trained STELLAR on the GVHD dataset using our gold-standard training set of 5k labeled cells, along with 150k unlabeled cells, and found that including the unlabeled data improved performance. The optimal model incorporated all proteins, constructed the spatial graph with a 20 μm distance threshold, and was trained for 12 epochs with a learning rate of 1e-3, weight decay of 5e-2, 23 attention heads, and a batch size of 32. The best-performing model identified three novel cell types, which were subsequently mapped to Neutrophil and two Fibroblast subclasses, based on the ground-truth test data to achieve the highest possible score.

#### CellSighter

CellSighter^28^ is a deep learning pipeline for automated cell classification in multiplexed imaging data, leveraging an ensemble of convolutional neural networks (CNNs) based on a ResNet50 backbone. The pipeline applies data augmentation during training, incorporates prediction confidence scores, and integrates multiple independently trained models into an ensemble for improved robustness. For benchmarking, we tested different batch sizes and hierarchical class structures. Next, we trained 10 independent models for 100 epochs, utilizing a batch size of 128, learning rate of 0.001, crop input size = 60, crop size = 128, including augmentation, and using the following 9 high-level cell classes: B cell, T cell, Myeloid, Mast, Plasma, Immune, Epithelial, VascFibro, Other. Model ensembles were created every 10 epochs by averaging the outputs of the trained models up to that point. The best results were obtained at 50 epochs. The final classification was determined by selecting the class with the top probability from the ensemble for each cell.

#### CELESTA

CELESTA^26^ is an unsupervised probabilistic method for cell classification in multiplexed imaging, combining marker-based and spatial-based classification to iteratively refine cell type assignments. The method first identifies high-confidence anchor cells based on their marker expression and then iteratively refines the classification of ambiguous cells using spatial neighborhood information. The approach requires a prior marker signature matrix defining marker expectations for each cell type, single-cell imaging data containing marker intensities and spatial information, and high and low expression thresholds to classify anchor and ambiguous cells. We tested multiple prior marker matrices, both with and without lineage categories, and found that excluding lineage information resulted in better classification performance for the GVHD dataset, as well as threshold and classification parameters. The final prior marker signature matrix included the following 33 cell types and their associated markers: B cell (CD20, CD45), CD4T (CD3, CD4, CD45), CD8T (CD3, CD8, CD45), Treg (CD3, CD4, FOXP3), Neutrophil (CD15, CD45), ImmuneOther_1 (CD11c, CD45), ImmuneOther_2 (CD11b, CD45), ImmuneOther_3 (HLADR, CD45), ImmuneOther_DC (CD11c, HLADR, CD45), ImmuneOther_APC (CD11c, CD11b, HLADR, CD45), ImmuneOther_CD45RA (CD45RA, CD45), Plasma (CD38, CD45), Endothelial (CD31, CD34, CD45), Fibroblast (SMA, PDPN, CD45), Epithelial (Cytokeratin, MUC1, NAKATPASE, CD45), Goblet (MUC1, CD45), Paneth (MUC1, CD45), Endocrine (SYP, CD45), Neuron (CD56, CD45), Neutrophil_CD15 (CD15, CD45), Plasma_CD38 (CD38, CD45), Fibroblast_1 (SMA, PDPN, CD45), Fibroblast_2 (Collagen, CD45), Fibroblast_Epithelial (Cytokeratin, SMA, CD45), Epithelial_1 (Cytokeratin, CD45), Epithelial_2 (Cytokeratin, MUC1, CD45), Macrophage_1 (CD68, CD163, CD45), Macrophage_2 (CD68, CD163, HLADR, CD45), Macrophage_3 (CD68, CD163, CD11b, CD45), Neutrophil_1 (CD15, CD11b, CD45), Neutrophil_2 (CD15, CD11b, HLADR, CD45), Neutrophil_CD15 (CD15, CD45), and Neuroendocrine (SYP, CD45). The best-performing threshold values were high_expr_thresh_anchor = (0.6, 0.5, 0.5, 0.1, 0.8, 0.5, 0.5, 0.1, 0.5, 0.5, 0.5, 0.5, 0.5, 0.3, 0.5, 0.5, 0.5, 0.5, 0.5, 0.5, 0.8, 0.1, 0.6, 0.7, 0.5, 0.1, 0.1, 0.8, 0.5, 0.5, 0.1, 0.1, 0.1), high_expr_thresh_index = high_expr_thresh_anchor - 0.1, and low_expr_thresh_anchor = low_expr_thresh_index = 1 for all cell types. For iterative refinement, we set max_iteration = 10 and cell_change_threshold = 0.01, meaning the algorithm stopped when fewer than 1% of cells changed classification between iterations. Cell types were grouped prior to comparison with our labels, e.g., Macrophage_1, Macrophage_2, and Macrophage_3 types were merged into Macrophage. Cells that did not meet the probability threshold for any category were classified as Unidentified.

#### Astir

ASTIR^30^ is a probabilistic model for cell type classification that integrates prior knowledge of marker proteins. We tested multiple marker file configurations, exploring different levels of resolution and hierarchical groupings. The final marker file that achieved the best results defined 20 cell types: B cell (CD20, CD45), CD4⁺T (CD3, CD4, CD45), CD8⁺T (CD3, CD8, CD45), Treg (CD3, CD4, FOXP3, CD45), CD3⁺T (CD3, CD45), ImmuneOther_1 (CD45, CD103, HLADRDPDQ), ImmuneOther_2 (CD45), Endocrine (CHGA, ECAD), Endothelial (CD31), Epithelial (ECAD, Keratin), Goblet (Mucin, ECAD), Fibroblast (SMA, Collagen), Macrophage_1 (CD68, CD163, CD206, CD14, CD45, HLADRDPDQ), Macrophage_2 (CD68, CD206, HLADRDPDQ), Mast (Tryptase, Lysozyme, ECAD, CD45), Neuron (CD56), Neutrophil (CD15, Calprotectin, Lysozyme, CD45, HLADRDPDQ), Paneth (Lysozyme, ECAD, Keratin), Plasma (IgA, CD38), and Unidentified. The hierarchical structure grouped these into broader categories: *vasc_fibro_cells* (Fibroblast, Endothelial, Neuron), *epithelial_cells* (Epithelial, Paneth, Goblet, Endocrine), and *immune_cells*, which included *other_immune* (ImmuneOther_1, ImmuneOther_2, Mast), *myeloid* (Neutrophil, Macrophage_1, Macrophage_2), and *lymphocytes* (plasma_b (Plasma, B cell), *tcells* (Treg, CD4⁺T, CD3⁺T, CD8⁺T)). Multiple hyperparameter configurations were also tested. The final settings used were max_epochs = 200, batch_size = 512, learning_rate = 0.003, n_init = 12, and n_init_epochs = 3. Cell type diagnostics confirmed that marker proteins were enriched in their respective cell types, and results were exported using a threshold of 0.7 for cell type assignment. Cell types were grouped prior to comparison with our labels, e.g., Macrophage_1 and Macrophage_2 types were merged into Macrophage.

#### Clustering

We used Self-Organizing Map (SOM) clustering^67^ to group cells based on their mean cell intensity for each protein. This is one of the most commonly used clustering methods for spatial proteomics data^68^. In addition to running on the intensities alone, SOM clustering was also applied for Nimbus, utilizing cell marker positivity probability features, and Pixie, utilizing cell pixel cluster features, as described in the following sections. We tested the Python implementation of SOM clustering from (https://github.com/angelolab/ark-analysis/tree/main/templates) used in the Nimbus^40^ and Pixie^25^ methods, as well as in R (FlowSOM^22^) and MATLAB^69^ (*selforgmap* function), with no observable differences between the implementations. Although the GVHD test/train data contains only 18 cell types, previous studies have demonstrated that overclustering cells into a larger number of groups before merging them into meta-clusters can improve classification performance^25^. To evaluate this approach, we tested 64 (8×8), 100 (10×10), 225 (15×15), 400 (20×20), and 900 (30×30) clusters, performing 10 replicates for each. We then applied hierarchical meta-clustering to 20, 32, 64, and 100 meta-clusters, provided the number of initial clusters exceeded the number of meta-clusters—e.g., for 64 clusters, only 20 and 32 meta-clusters were tested. To establish an upper bound on classification performance, we assigned both clusters and meta-clusters to cell types using the ground-truth test set with a majority vote approach: ground truth labels were evaluated for all cells in the cluster, which subsequently received the label pertaining to the majority. However, for the configuration with 900 initial clusters, we only assigned labels at the meta-cluster level rather than to individual clusters, as manually inspecting and labeling 900 separate clusters would be prohibitively time-consuming in any practical workflow. We then refined the cluster assignments to optimize the F1 score, ensuring that rare cell types were not misclassified into more abundant ones, as missing these rarer populations had a greater negative impact on the overall score. Across all configurations, the best results were achieved using 400 clusters, directly assigned to cell types without meta-clustering.

#### Nimbus

Nimbus^40^ is a deep learning model designed to classify marker positivity (+/-) from multiplexed imaging data by processing image tiles and segmentation masks. It utilizes a PanopticNet-based architecture, which combines semantic segmentation and instance segmentation to extract features from individual cells. The model applies post-processing steps such as test-time augmentation, tile-and-stitch inference, and confidence score integration to refine classifications. Marker intensities were normalized to the 99.9th percentile, with low-expressing markers (e.g., CD20) adjusted to the 99.99th percentile. The normalization factors were: CD4 (5.40), CD20 (0.78), CHGA (10.49), FOXP3 (1.05), Mucin (6.82), CD14 (3.27), CD69 (3.23), IDO1 (0.97), Keratin (20.93), CD68 (4.66), CD103 (2.23), CD3 (2.83), NAKATPASE (5.21), CD31 (4.77), Ki67 (4.53), GZMB (3.37), VIM (8.00), IgA (2.60), CD45RO (4.00), Calprotectin (2.67), Tryptase (20.78), TCF (3.21), CD56 (4.60), CD45RA (3.01), PD1 (0.05), CD206 (3.83), CD209 (2.22), SMA (101.19), PDL1 (0.66), CD8 (1.98), CD45 (9.48), HLA1 (8.54), CD38 (6.68), dsDNA (84.04), Collagen (29.54), ECAD (7.64), HLADRDPDQ (11.39), CD15 (4.06), CD163 (2.94), Lysozyme (4.44). We applied SOM clustering, as described above, to cluster the cells based on the marker positivity probabilities predicted by Nimbus.

#### Pixie

Pixie^25^ is a pixel clustering-based approach for analyzing multiplexed imaging data, utilizing Self-Organizing Maps (SOMs) to group pixels and then cells based on their marker expression profiles. Pixie first performs pixel-level classification, followed by cell-level aggregation, and hierarchical meta-clustering. For this analysis on GVHD, pixel clustering was performed on 30 protein markers: CD103, CD14, CD15, CD163, CD20, CD206, CD3, CD31, CD38, CD4, CD45, CD45RA, CD56, CD68, CD8, FOXP3, HLADRDPDQ, IgA, Calprotectin, ECAD, Keratin, VIM, Collagen, Mucin, SMA, NAKATPASE, CHGA, Lysozyme, Tryptase, and dsDNA. Prior to clustering, a Gaussian blur with a factor of 1 was applied to smooth the imaging data, and normalization was performed such that each pixel’s intensity was scaled by the sum of all channels. To optimize computational efficiency, 50% of pixels per field of view (FOV) were randomly sampled for training. We tested different parameters including the numbers of pixel clusters and number of pixel metaclusters ranging from 64 to 900. Ultimately the best performance was pixel clustering with 400 pixel clusters and 300 meta-clusters after applying z-score scaling and capping at a value of 3. To assign clusters to biologically meaningful cell types, we tested both automatic classification using ground-truth labels and manual curation (manually grouping pixel clusters based on their expression), which required substantial time. In each case cell clustering was performed as described above using the aggregated pixel cluster values with and without the intensity values. A completely independent test by a different researcher on a smaller subset of the data (33,000 cells from 18 FOVs) confirmed that the ultimate cell classification accuracy remained consistent across different parameter settings and different users.

#### Clustering & Curation

A common approach to improving classification accuracy in multiplexed imaging studies is to cluster cells based on their expression profiles and then manually curate the labels by correcting misclassifications. Since no automated method achieves perfect accuracy, this process involves overlaying key markers on the image and systematically refining classifications. For example, by visualizing CD8 expression, a researcher can correct CD8^+^T cell classifications—both by relabeling missed classifications (cells expressing CD8 but assigned to a different cluster) and by identifying incorrect classifications (cells lacking CD8 expression but erroneously labeled as CD8+ T cells). The latter is particularly challenging, as it requires time to relabel the cell to one of the other 17 cell classifications. This curation process is highly labor-intensive and becomes impractical for achieving high accuracy across very large datasets. Still, prior to the development of CellTune, this manual curation approach—combining clustering-based classification with expert correction—was our method of choice for achieving the highest possible accuracy in multiplexed imaging data. To assess the impact of expert curation, one expert labeler performed a detailed manual refinement of a subset of the GVHD dataset (33,000 cells across 18 FOVs) that had been initially classified using Pixie’s pixel clustering and cell classification. The expert labeler spent over 80 hours reviewing classifications on a cell type–by–cell type basis, systematically making corrections to improve accuracy.

#### Optimized Gating

Gating, a technique originally developed for single-cell methods, is widely used for classifying cells based on marker expression thresholds. To establish an optimized gating approach for multiplexed imaging data, we performed both manual gating and an semi-automated “optimized” gating classification using simple decision trees. The decision trees were trained to classify one cell type at a time based on the test data in MATLAB^69^ with the *fitctree* function, limiting the number of splits (<20) and manually adjusting/reusing many gates on individual channels (e.g. CD45 > or < 4 determining positive / negative) to more closely mimic manual gating. At each step, classification was only performed on the remaining unclassified cells. We tried different ordering of cell types, and the best results were obtained by starting with the rarest cell types, such as CD3T and Treg, and progressively classifying more abundant populations, finishing with Fibroblast and Epithelial cells. In all cases, the decision tree-derived gates outperformed our tests of manual gating, thus defining an upper bound for optimized gating performance. The final optimized gates are shown in parenthesis for each cell type as follows: NK (CD56 > 3.9, GZMB > 2.8, CD45 > 4, CD3 < 3.5, HLADRDPDQ > 4.4); CD3T ((CD4 < 2 & CD8 < 2) & ((CD3 > 3.8 & HLADRDPDQ < 5.7) | (CD3 > 3 & HLADRDPDQ < 4.3))); Treg (FOXP3 > 4); Endocrine (CHGA > 5.5); Mast (Tryptase > 5.5 & CD206 < 3 & CD68 < 3.5 & CD163 < 4.5 & SMA < 6); Paneth_1 (Lysozyme > 4 & Keratin > 3 & Mucin < 3); Neutrophil (Calprotectin > 4 | (Calprotectin > 3 & (CD15 > 4 & CD68 > 3)) | (CD15 > 4 & Keratin < 3 & Mucin < 3)); Endothelial (CD31 > 4 & SMA < 6 & CD45 < 4 & IgA < 3.5 & CD206 < 3 & CD68 < 3.5 & CD163 < 4.5); B cell (CD20 > 3 & CD45RA > 5 & CD3 < 4); Neuron (CD56 > 4 & CD45 < 4 & IgA < 3.5 & CD206 < 3 & CD68 < 3.5 & CD163 < 4.5); CD8T ((CD206 < 3 & CD163 < 4.5 & CD68 < 3.5) & (CD8 > 4.5 | (CD8 > 3.5 & CD4 < 4.5) | (CD8 > 2.5 & CD4 < 4.5 & (GZMB > 3 | CD3 > 3))); Goblet (Mucin > 5 | (Mucin > 4 & ECAD < 4 & NAKATPASE < 3.5 & CD45 < 4)); CD4T (CD3 > 3 & CD4 > 0.5 & CD206 < 3, CD68 < 3.5 & CD163 < 4.5 & SMA < 6); Plasma cells (CD206 < 3 & SMA < 6 & CD68 < 3.5 & CD163 < 4.5 & (IgA > 3.5 | (Keratin < 3 & CD38 > 3.5 & CD68 < 2))); Macrophage ((CD206 > 3 & SMA < 6 & Keratin < 3) | (CD68 > 3.5 & SMA < 6 & Keratin < 3) | (CD14 > 2.5 & SMA < 6 & Keratin < 3) | (CD163 > 4.5 & SMA < 6 & Keratin < 3)); Fibroblast (SMA > 6 | (Collagen > 4 & CD45 < 4 & Keratin < 3)); Epithelial (CD45 < 4 & ((Keratin > 3 & VIM < 3) | ((Keratin > 3 | VIM < 3) & (ECAD + NAKATPASE) > 3 & SMA < 6 & Lysozyme < 4))); Paneth_2 (CD45 < 4 & Lysozyme > 4 & (VIM < 3 | (Keratin < 3 & (ECAD + NAKATPASE) > 3))); ImmuneOther (CD45 > 4 & SMA < 6); The remaining cells were labeled as Unidentified. These thresholds were applied uniformly across the dataset. While additional improvements could be achieved by gating each image individually, this would require a significant time investment. This optimized gating strategy relied on ground-truth labels, and thus serves as an upper bound for the accuracy that can be achieved by gating.

#### Ensemble

For comparison, we also constructed a majority vote ensemble based on the predictions of nine of the methods above – excluding Clustering & Curation, which was performed on only a subset of the data. Each cell was assigned the label receiving the highest number of votes among the nine methods (range: 3-9 votes). In cases of ties (e.g., 3-3 or 4-4 votes between labels), we assigned the label matching the consensus ground truth, ensuring a fair evaluation.

### Evaluation of Label and Feature Importance

#### Simulations of the CellTune classification workflow

To evaluate how different aspects of the CellTune training process impact classification performance— including transfer learning to novel datasets, landmark initialization, and sampling strategies for labeling— we conducted simulations of the CellTune workflow. In all cases, classifiers were first initialized with landmark-labeled cells, representing the starting point of the simulation (i.e., zero added labels on the x-axis). From this baseline, we simulated multiple labeling cycles in which additional labels were introduced, a new model was trained, and predictions were evaluated.

For simulations assessing transfer learning and landmark initialization strategies (**Fig. 4B,H**), we used manual labels generated through actual CellTune annotation sessions. In the transfer learning simulations (**Fig. 4B**), classifiers were initialized with gated landmark cells from the GVHD dataset and supplemented at initialization with training labels from PDAC, NSCLC, and/or melanoma datasets. Additional labels were then drawn from the GVHD dataset during subsequent training cycles using CellTune’s query-by-committee sampling strategy with default parameters. In the landmark initialization simulations (**Fig. 4H**), all training labels were from the GVHD dataset, but the initial landmark cells were selected either manually or using automated landmarking based on marker positivity predictions. In both cases, performance was evaluated on the gold-standard consensus labeled test set for GVHD.

For simulations evaluating label selection strategies (**Fig. 5A**), we used CellTuneDepot’s high-confidence labels as ground truth for both sampling training labels and evaluating predictions. This approach enabled systematic comparison of multiple labeling strategies—random sampling, by field of view (FOV), and CellTune’s query-by-committee (QBC)—without requiring new manual annotations for each simulation. Generating a dedicated set of manual labels for each strategy would be impossible, requiring a prohibitive time investment in manual labeling. To ensure a fair comparison, all simulations used the same pool of high-confidence labels, the same evaluation set, and the same initial set of landmark-labeled cells; only the sampling strategy used to select additional training labels varied.

#### Feature ablation analysis

An ablation study was performed to assess the contribution of individual features to the performance of the CellTune classifier. Since the full model was trained on 1802 features, to make the analysis more informative, we grouped the different features into groups of related features. As detailed in the “Compute features” section of the Methods, each feature is constructed as a combination of four components: a channel, a statistical measurement, a spatial location, and a cell assignment, i.e., self or neighbor (**Fig. S1B**). For the ablation study, the feature groups were defined as subsets of the space [{Channels}, {Statistics}, {Locations}, {Cells}], where * denotes all possible values within an element. The *Baseline* group consists of the standard mean expression features within each cell. The order of the remaining feature groups was determined by the difficulty in generating the features, from the easiest (Max) to the most computationally difficult and/or time consuming (Subregion). Features that could belong to multiple groups went to the most difficult one; for example, a feature combining *Subregion*, *Max*, *Erode*, and *Neighbor* components was assigned to the *Subregion* group rather than the *Max*, *Erode*, or *Neighbor* groups. The specific groups were defined as follows: *Baseline* (*Mean*) *=* [{Proteins}, {Mean}, {Cell}, {Self}]; *Max* = [{Protein}, {Max}, {Cell}, {Self}]; *Var* = [{Protein}, {Var}, {Cell}, {Self}]; *Neighbor* = [{Protein}, {Mean, Max, Var}, {Cell}, {Neighbor}]; *Composite* = [{Composite}, {Mean, Max, Var}, {Cell}, *]; *Erode* = [{Protein, Composite}, {Mean, Max, Var}, {Erode}, *]; *Dilate* = [{Protein, Composite}, {Mean, Max, Var}, {Dilate}, *]; *Environment* = [{Protein, Composite}, {Mean, Max, Var}, {Environment}, *]; *Counts* = [{Protein, Composite}, {Counts}, *, *]; *Colocalization* = [{Colocalization}, *, *, *]; *Subregion* = [{Subregion}, *, *, *]; *Full Model* = [*, *, *, *]; (Note the marker positivity probability features were excluded from the ablation study.) For each configuration, a CellTune classifier was trained with GVHD manual labels, using only the specific subset of features from the corresponding group(s).

The ablation study was performed in three regimes: *Add*, *Remove* and *Accumulate*. *Add* started with a baseline model, which included only the feature of the mean cellular expression per protein, adding one feature group at a time. This configuration did not account for interactions or compensations between feature groups. For example, to assess the contribution of pixel counts (Counts), the model was trained using only the baseline features and the pixel counts per cell. In this setup, a higher F1 score indicates greater importance of the added feature group. In the *Remove* configuration, the model was trained using all features except the feature group being evaluated. For instance, to determine the impact of the Counts feature group, the model was trained on all features except pixel counts. Unlike the *Add* configuration, this approach allowed assessment of interactions between feature groups. In this setup, a lower F1 score indicates greater feature importance, as its removal negatively impacted classifier performance. In the *Accumulate* configuration, models were trained by sequentially adding feature groups to the baseline model. The order of addition was dictated by the difficulty in generating the features, from the easiest (Max/Var) to the most computationally difficult and/or time consuming (Subregion). In this setup, a higher increase in F1 score indicates greater feature importance.

We observed that some features can compensate for others. For example, in the Goblet cells ablation study (**Fig. S4**), the *Add* configuration shows that incorporating spatial features improves model performance. However, the *Remove* analysis does not highlight these features. This discrepancy arises because certain features can act as substitutes for one another, allowing the model to adjust its decision boundaries by finding alternative feature splits that combine or separate features in different ways to compensate for the missing information.

#### SHAP feature importance

SHAP values were calculated using the SHAP Python package (v.0.42.1) to assess the importance and contribution of different features to the classification. A TreeExplainer object was employed to compute SHAP values for both summary and waterfall plots. For the summary plot of the XGBoost model, SHAP values were computed on all cells from the GVHD dataset, while for the CatBoost model, SHAP values were calculated on 100K cells due to memory limitations.

### Statistics and Figures

All statistics were computed using Python^70^ (3.9). Plots were generated using MATLAB^69^ (R2022b) and the Matplotlib^71^ (3.9) and seaborn^72^ (0.13) libraries in Python. Figures were prepared using Biorender^73^ and Adobe Illustrator. For visualization purposes only, images were clipped to their dynamic range. Some MIBI images were Gaussian blurred using the SciPy^74^ library in Python.

### Software

CellTune (0.1) is implemented in Python^70^ (3.9) with a GUI built using PyQt^75^ (5.15) and superqt^76^ (0.6). Image processing and data analysis utilize OpenCV^77^ (4.9), scikit-image^78^ (0.24), SciPy^74^ (1.13), NumPy^79^ (1.26), pandas^80^ (2.2), and Matplotlib^71^ (3.9). Machine learning models are supported via scikit-learn^81^ (1.5), XGBoost^82^ (2.0), CatBoost^45^ (1.2), and LightGBM^83^ (4.5), while deep learning models use PyTorch^66^ (2.6). Database management is handled with SQLAlchemy^84^ (2.0). The software is packaged using PyInstaller^85^ (6.7).

## Data Availability

Data is available through the CellTuneDepot repository at https://celltune.org/celltunedepot. CellTuneDepot is divided into sections for high-quality labels and manual labels, and further organized by dataset. Each dataset includes multiplexed images, segmentations, channel and cell type information, populations labels, and a cell table containing the mean expression of each protein for each cell.

## Software Availability

CellTune software is available as a standalone desktop application at https://celltune.org.

## Acknowledgements

We would like to thank Lacey Padrón and Robin Kageyama of the Parker Institute for Cancer Immunotherapy for their support with the Mantis Viewer software used for testing CellTune’s classification method in its early stages of development, Reinat Nevo for valuable discussions, Vitaly Golodnitsky for IT assistance, and the Department of Life Sciences Core Facilities of the Weizmann Institute of Science for providing access to computational resources. L.K. holds the Fred and Andrea Fallek President’s Development Chair and is supported by the Enoch foundation research fund, the Abisch-Frenkel foundation, the Rising Tide foundation, the Sharon Levine Foundation, Fundación Alberto Palatchi, Dwek center for cancer immunotherapy, and grants from the European Research Council (948811), Israel Science Foundation (2481/20), the Rosetrees Foundation (10004), the Israel Precision Medicine Partnership Program (3830/21), and the Melanoma Research Alliance Team Science Award (1200724). Y.B. is supported by the Clore Scholars Program of the Clore Israel Foundation.

---

**Figure S1:**
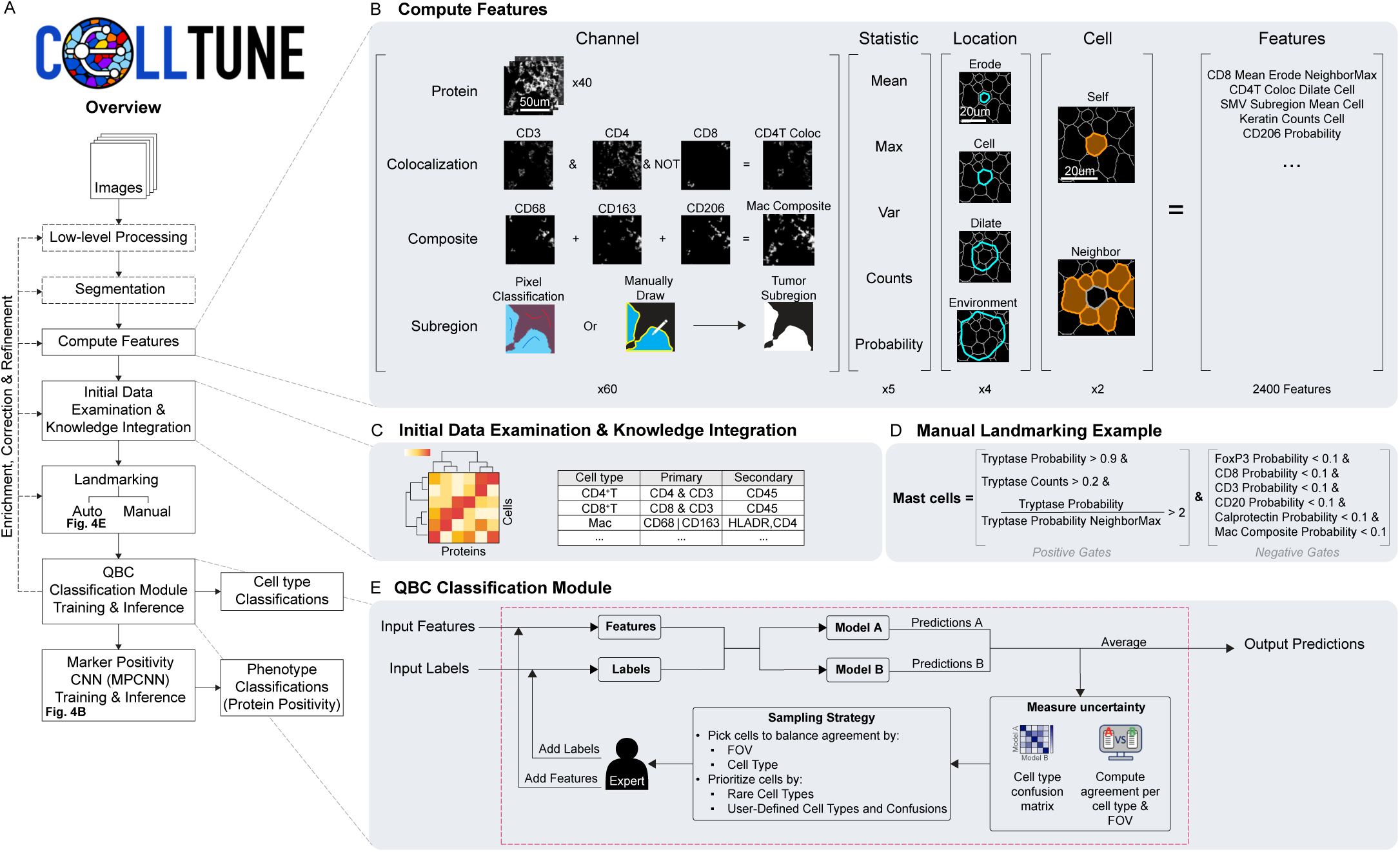
Detailed schematic of CellTune’s cell classification workflow. **(A)** Overview of the complete pipeline from protein images to final cell type and phenotype (protein positivity) classifications. The segmentation step is not integrated into the first version of CellTune but will be in future updates. **(B)** Comprehensive cell feature computation performed in CellTune. Each feature is derived from a channel, including single protein expression, multiple protein colocalization, composites of proteins, and subregions (via pixel classification or manual drawing). Four statistical measures (mean, maximum, variance, and pixel count) are computed across distinct cellular compartments: the cell center (erode), entire cell segmentation (cell), immediate surroundings (dilate), and broader cellular environment (environment). Features represent either an individual cell’s measurements or the maximum value among its neighboring cells. In total, approximately 2000 features are computed per cell, depending on the number of channels. **(C)** Prior to landmarking and classification, initial data examination is conducted, in which the user integrates prior knowledge with data visualization to identify known cell types and their associated lineage proteins, as well as to determine if any additional colocalization, composite, or subregion channels are needed. **(D)** Manual landmarking example of mast cells, applying a series of positive and negative thresholds. Positive gates are placed on the primary lineage channel(s) for the cell type. Negative gates are applied to exclude cells with high values for the primary markers of other cell types. Users can easily adjust thresholds, adding or removing gates according to their specific needs. Alternatively, users can use CellTune’s automated landmarking method depicted in Figure 4E. **(E)** The Query-by-Committee (QBC) classification process begins with initial cell labels and computed features. In each iteration, two independent models are trained using these labels and features. Uncertainty is assessed by evaluating the agreement between the two models’ predictions, summarized overall, per cell type, and per field-of-view (FOV), generating a confusion matrix highlighting pairwise disagreements across the dataset. Cells with conflicting predictions are strategically sampled, balancing agreement across FOVs and cell types, while prioritizing rare and user-specified cell types and cell type confusions. The user reviews and labels the sampled cells, augmenting the training set. During labeling, users can introduce new cell types and define additional features. Importantly, this interactive labeling step can reveal upstream issues with low-level processing and segmentation, enabling early troubleshooting before extensive downstream analysis. Iterations (or cycles) continue until classification results are satisfactory. Final predictions from both models are averaged. **(A: MPCNN Training & Inference)** Phenotype classifications are based on marker positivity of functional proteins. Marker positivity is calculated by running the inference step of the Marker Positivity Convolutional Neural Network (MPCNN), as depicted in Figure 4B. An MPCNN which was trained on CellTuneDepot is supplied with CellTune. Conversely, the user may choose to train a dataset-specific MPCNN as depicted in Figure 4B. Both training and inference are accessible via the CellTune software.

**Figure S2:**
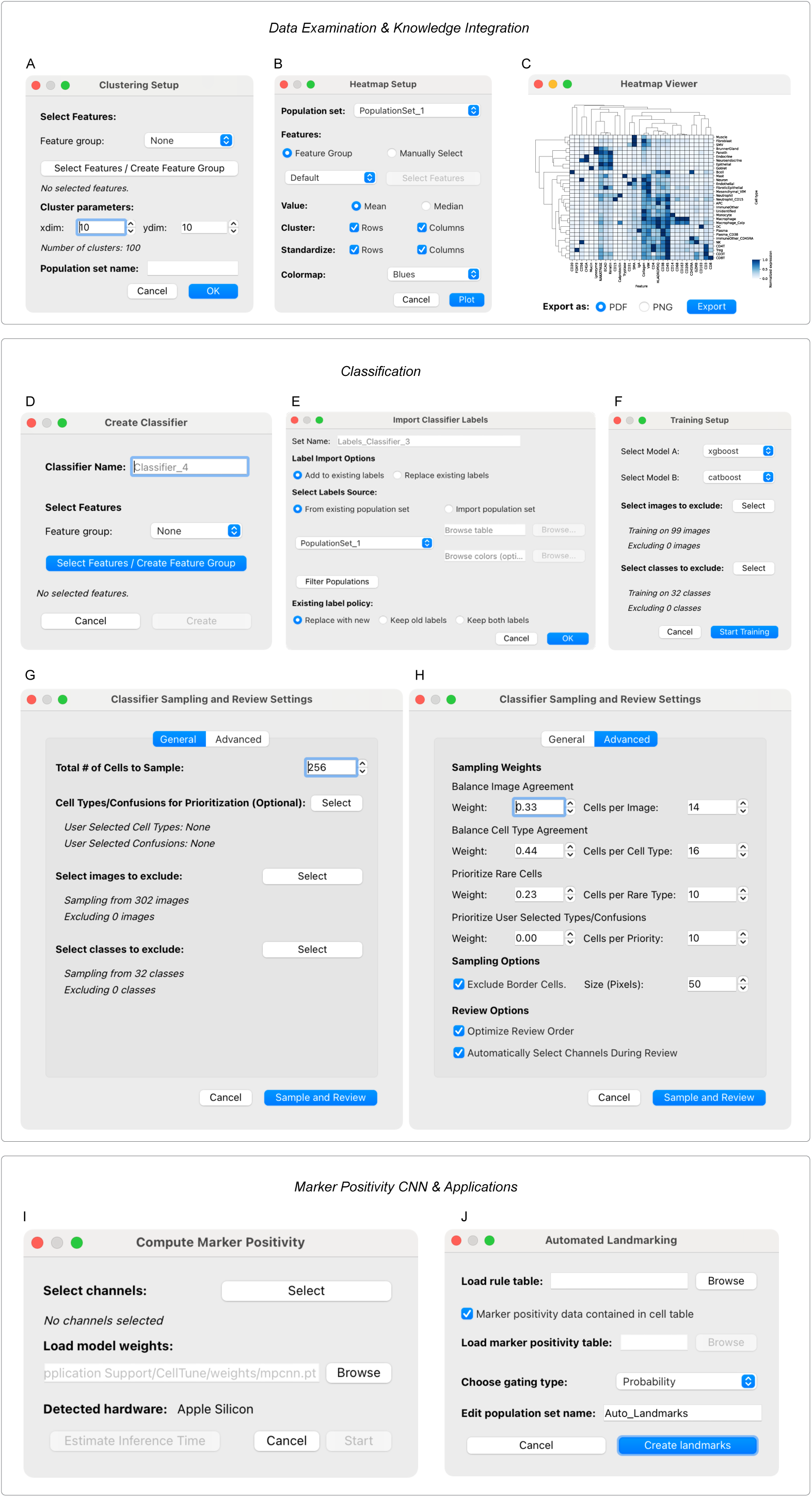
The CellTune software provides comprehensive interfaces supporting an integrated analysis pipeline, including data exploration, classification, and marker positivity inference. **(A)** Clustering for initial data examination is performed using the *Clustering Setup* dialog. The user selects the features for clustering and specifies the number of clusters. A population set is created from the resulting clusters. **(B)** Data visualization via heatmap is enabled from the *Heatmap Setup* dialog. The user selects the population set and features to visualize, along with other heatmap settings. **(C)** Heatmap viewer in the software. **(D-H)** The software facilitates cell classification functionalities. **(D)** The user creates a cell classifier and selects the features used for classification. **(E)** Labels are imported into the classifier for training, with the option to add new labels during the process. **(F)** The *Training Setup* dialog allows users to choose model architectures and specify images or classes to exclude from training. **(G-H)** Settings for sampling and review after training. **(G)** Users control the number of sampled cells after a training cycle, prioritize specific cell types or confusions, and define exclusions for images or classes. **(H)** Advanced settings allow fine-tuning of the sampling strategy, including prioritization and balancing. **(I)** Computation of marker positivity using MPCNN. Users select the channels for which marker positivity probabilities will be computed. **(J)** The *Automated Landmarking* dialog allows users to load a landmarking rule table to automatically create landmark cells.

**Figure S3:**
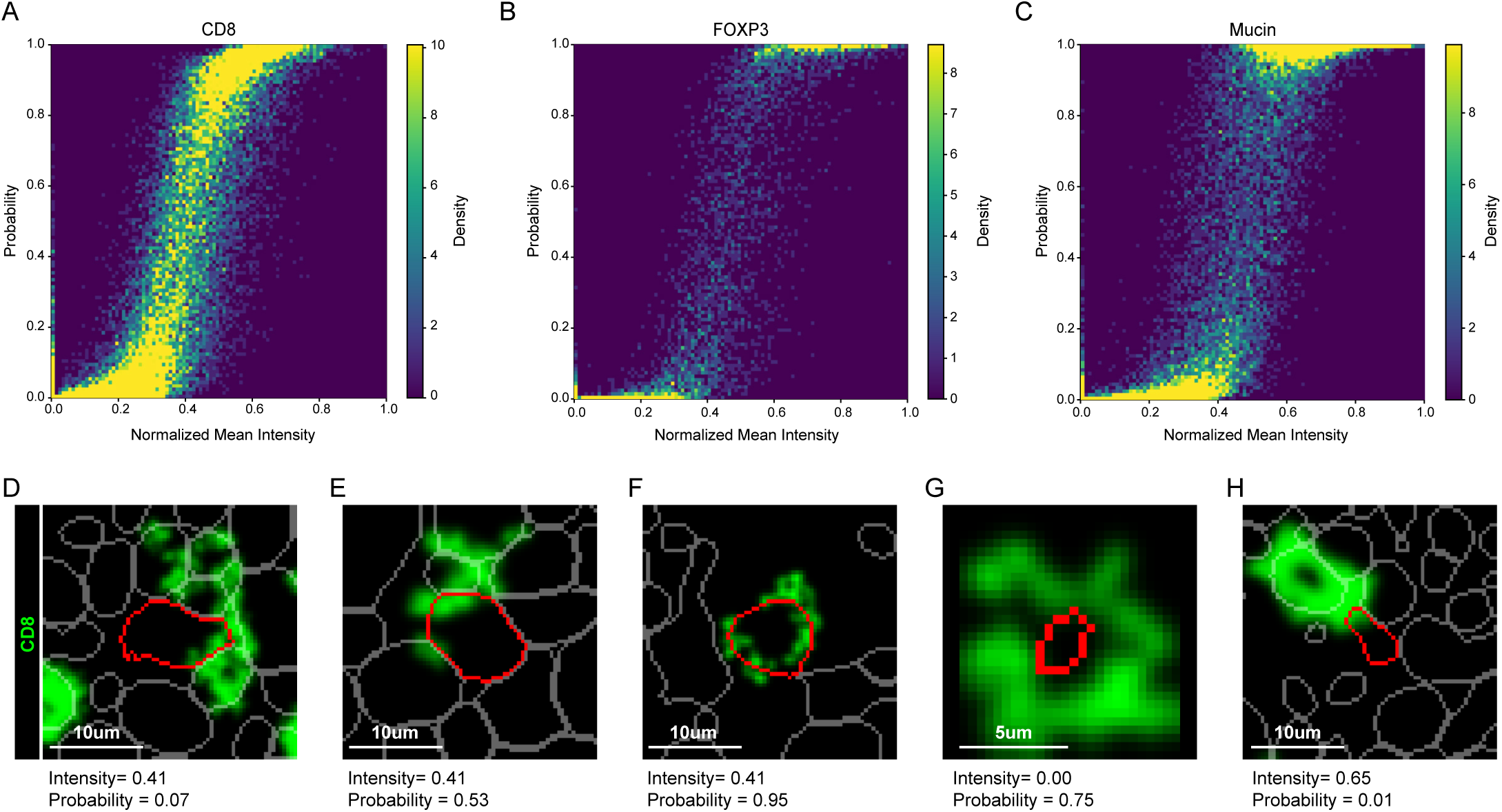
MPCNN probabilities exhibit variability beyond protein intensity. Density heatmaps depicting the relationship between MPCNN-generated marker positivity probabilities and normalized mean protein intensities for **(A)** CD8 (membranous), **(B)** FOXP3 (nuclear), and **(C)** Mucin (cytoplasmic). While probabilities generally correlate with protein expression, cells with intermediate intensity exhibit a wide range of predicted probabilities, particularly pronounced for membranous markers like CD8 due to signal spillover **(D-F)**. Examples of cells with the same normalized mean CD8 intensity but having low **(D)**, medium **(E)**, and high **(F)** CD8 probabilities. **(G)** Some cells having zero intensity and high probability occur due to signal missed by segmentation. **(H)** Conversely, a cell may have high normalized mean intensity due to signal spillover and the network correctly predicts it has low positivity probability.

**Figure S4:**
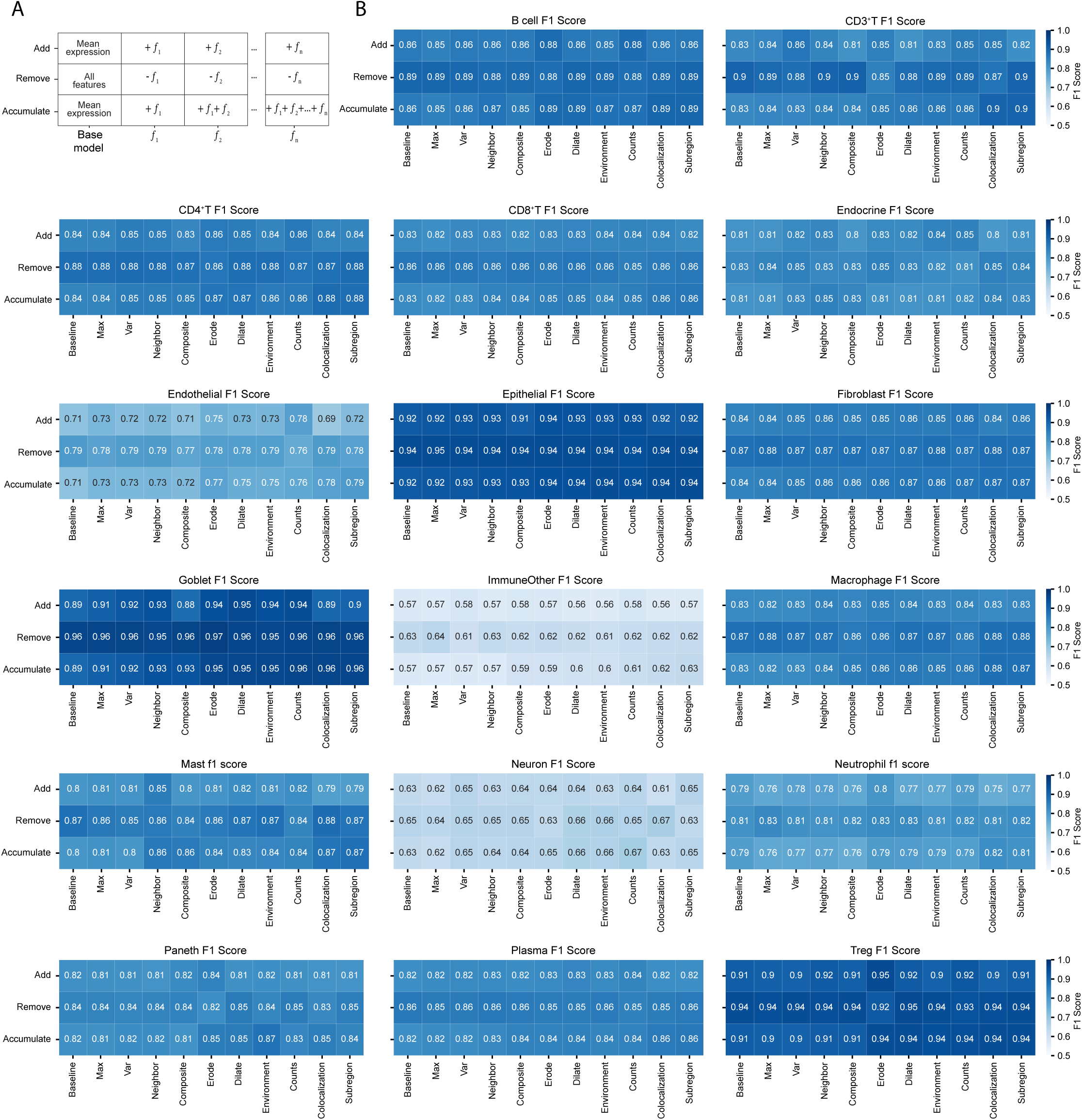
**Ablation study results across GVHD cell types**. **(A)** Scheme of the ablation study. All 1802 features were grouped into related feature types. CellTune simulations were conducted with different feature types. *Add:* adding a single feature type to the baseline model. *Remove:* removing a single feature type from the full model. *Accumulate:* adding and accumulating feature types to the baseline model. Features are added in order of increasing complexity to generate them. **(B)** Ablation study F1 scores per cell type.

