## Supplemental Table 1 for "CellTune: An integrative software for accurate cell classification in spatial proteomics"

**Supplementary Table 1: Number of cell types in published datasets**

| Method | Dataset ID | Imaging Platform | Tissue | Cell Types | Dataset Publication |
|---|---|---|---|---|---|
| Astir | Astir_DS1 | IMC | Breast | 4 | Schapiro et al. 2017 |
| Astir | Astir_DS2 | CyTOF | Breast | 6 | Wagner et al. 2019 |
| Astir | Astir_DS3 | CyCIF | Multi | 7 | Lin et al. 2018 |
| Astir | Astir_DS4 | IMC | Breast | 8 | Jackson et al. 2020 |
| CELESTA | Celesta_DS1 | CODEX | HNSCC | 8 | Zhang et al. 2022 |
| CellSighter | CellSighter_DS1 | MIBI | GI | 8 | Milo et al. 2024 |
| MAPS | MAPS_DS1 | MIBI | DLBCL | 9 | Wright et al. 2023 |
| CellSighter | CellSighter_DS2 | MIBI | IMC | 9 | Hoch et al. 2022 |
| CellSighter | CellSighter_DS3 | MIBI | Melanoma | 10 | Keren *Unpublished* |
| STELLAR | STELLAR_DS1 | CODEX | Tonsil | 10 | Hickey 2022 |
| MAPS | MAPS_DS2 | MIBI | cHL | 12 | Shaban et al. 2024 |
| CelloType | CelloType_DS1 | CODEX | CRC | 12 | Schürch et al. 2020 |
| CELESTA | Celesta_DS2 | CODEX | CRC | 13 | Schürch et al. 2020 |
| MAPS | MAPS_DS3 | MIBI | cHL | 13 | Shaban et al. 2024 |
| CellSighter | CellSighter_DS4 | CODEX | CRC | 13 | Schürch et al. 2020 |
| Pixie | Pixie_DS1 | MIBI | LN | 13 | Liu et al. 2023 |
| STELLAR | STELLAR_DS2 | CODEX | Barrett's Esophagus | 13 | Hickey 2022 |
| CelloType | CelloType_DS2 | CODEX | Bone Marrow | 13 | Bandyopadhyay et al. 2024 |
| MAPS | MAPS_DS4 | CODEX | CRC | 14 | Schürch et al. 2020 |
| CellSighter | CellSighter_DS5 | MIBI | Melanoma_LN | 14 | Keren *Unpublished* |
| CellTune_Manual | CellTune_Manual_DS1 | CODEX | Breast | 14 | Ben-Uri et al. 2025 |
| MAPS | MAPS_DS5 | CODEX | cHL | 16 | Shaban et al. 2024 |
| Pixie | Pixie_DS2 | MIBI | TNBC | 17 | Liu et al. 2023 |
| CellTune_Manual | CellTune_Manual_DS2 | CODEX | CRC | 17 | Schürch et al. 2020 |
| CellTune | CellTune_DS1 | CODEX | CRC | 17 | Schürch et al. 2020 |
| CellTune_Manual | CellTune_Manual_DS3 | MIBI | GI | 18 | Milo et al. 2024 |
| CellTune_Manual | CellTune_Manual_DS4 | MIBI | Melanoma | 20 | Keren *Unpublished* |
| CellTune | CellTune_DS2 | MIBI | Melanoma | 20 | Keren *Unpublished* |
| STELLAR | STELLAR_DS3 | CODEX | GI | 21 | Hickey et al. 2021 |
| CellTune | CellTune_DS3 | MIBI | PDAC | 27 | Keren *Unpublished* |
| CellTune | CellTune_DS4 | MIBI | Vlelanoma Brain Mets | 29 | Keren *Unpublished* |
| CellTune | CellTune_DS5 | MIBI | NSCLC | 30 | Keren *Unpublished* |
| CellTune | CellTune_DS6 | MIBI | GI | 32 | Milo et al. 2024 |
